# The membrane-targeting-sequence motif is required for exhibition of recessive resurrection in *Escherichia coli* RNase E

**DOI:** 10.1101/2024.04.06.588376

**Authors:** Papri Basak, Manjula Ekka, Apuratha Pandiyan, Smriti Tandon, Jayaraman Gowrishankar

## Abstract

The essential homo-tetrameric endoribonuclease RNase E of *Escherichia coli* participates in global RNA turnover as well as stable RNA maturation. The protomer’s N-terminal half (residues 1-529) bears the catalytic, allosteric and tetramerization domains, including the critical active site residues D303 and D346. The C-terminal half (CTH, residues 530-1061) is dispensable for viability. We have previously described a phenomenon of recessive resurrection in RNase E that requires the CTH, wherein the wild-type homo-tetramer apparently displays nearly identical activity in vivo as a hetero-tetramer comprised of three catalytically dead subunits (with D303A/D346A substitutions) and one wild-type subunit. Here we show that recessive resurrection is exhibited even in dimeric RNase E with the CTH, and that it is largely dependent on presence of the membrane-targeting-sequence motif (residues 565-582). A single F575E substitution also impaired recessive resurrection, whereas other CTH motifs (such as those for binding of RNA or of partner proteins) were dispensable. The phenomenon was independent of RNA 5’-monophosphate sensing by the enzyme. We propose that membrane-anchoring of RNase E renders it processive for endoribonucleolytic action, and that recessive resurrection and dominant negativity associated with mutant protomers are mutually exclusive manifestations of, respectively, processive and distributive catalytic mechanisms in a homo-oligomeric enzyme.

## Introduction

The essential endoribonuclease RNase E in *Escherichia coli* and *Salmonella enterica* not only catalyzes the rate-limiting initial steps in global RNA degradation and turnover (1) but also participates in gene regulation mediated by small RNAs as well as in rRNA and tRNA processing and maturation reactions (2–12). It is a large homo-tetrameric enzyme whose subunit polypeptides (encoded by the *rne* gene) are each comprised of 1061 amino acid residues. Intracellular activity of RNase E is autoregulated, since the 5’-untranslated region of *rne* mRNA is itself a substrate for cleavage by the enzyme (13–19).

Several features of RNase E are depicted in Figure 1A (top line) on a linear polypeptide representation. The N-terminal half of RNase E (NTH, residues 1 to 529) is by itself sufficient for homo-tetramer assembly as well as to confer bacterial viability. Crystal structures have been determined of the RNase E-NTH without and with bound RNA (20–23). Each subunit is folded into a large domain (residues 1 to 400) and a small domain (residues 438 to 529), with a pair of Cys residues (C404, C407) in the intervening linker region. The large domain comprises the catalytic active site that includes residues D303 and D346 as well as an allosteric regulatory site described further below. The enzyme exists as a dimer of dimers, with a Zn^2+^ ion co-ordinated between the Cys residues of a pair of linker regions contributing to dimer formation (referred to as a principal dimer); inter-subunit contacts between the pair of small domains of one principal dimer with those of a second then results in tetramer assembly.

**Figure 1:**
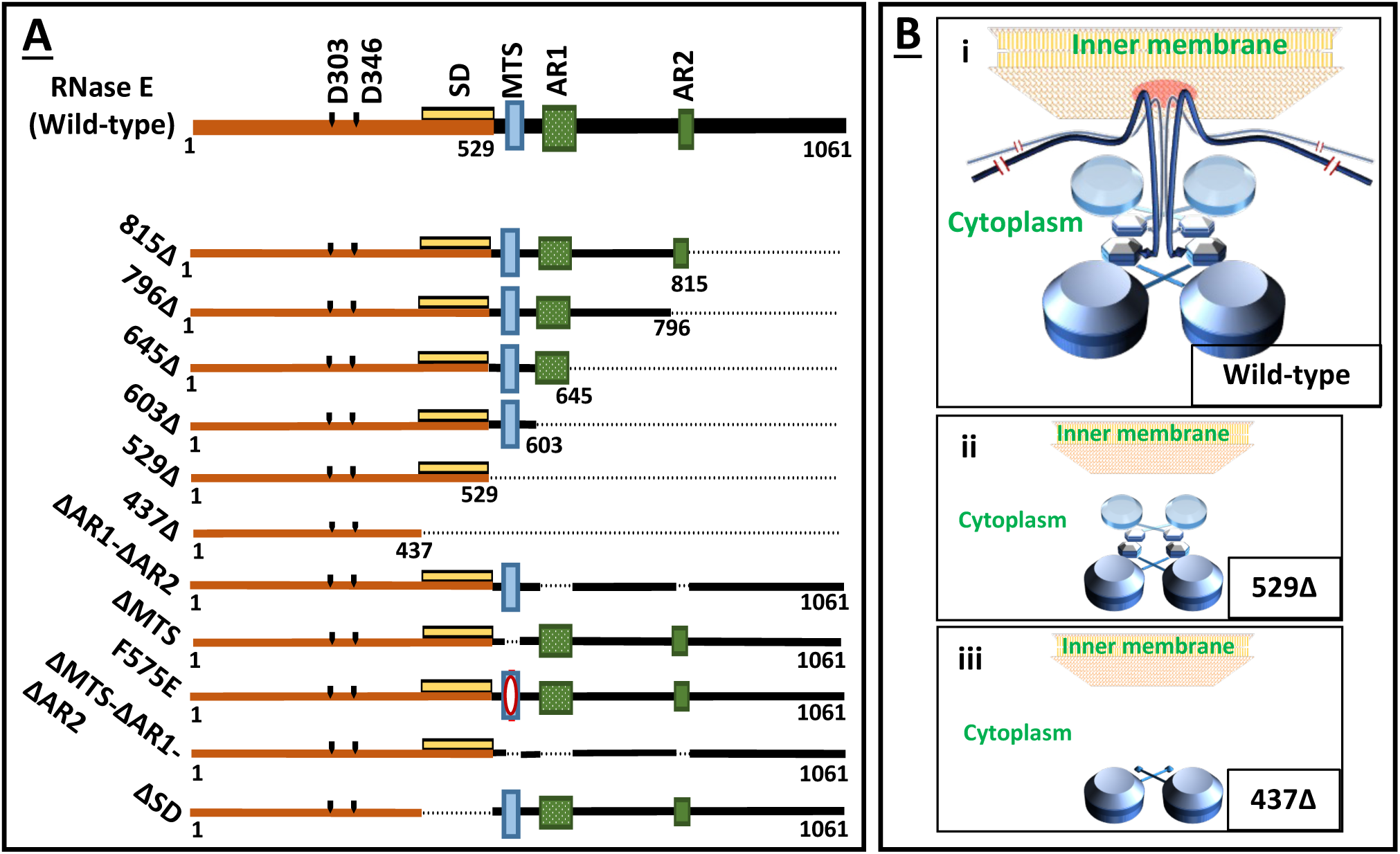
Features of RNase E and its variants. (**A**) Presented on top is a linear representation (not to scale) of the 1061-amino acid long wild-type RNase E polypeptide, with the positions of different motifs and active site residues marked; the NTH and CTH regions are shown in orange and black, respectively. Sites for binding of other proteins are not shown. Each of the lines beneath represents an ASKA plasmid-encoded RNase E variant used in this study; deletions (in-frame or at the C-terminus) are denoted by interrupted lines. (**B**) Schematic depiction (not to scale) of wild-type RNase E (panel i) and of its truncation variants 529∆ (panel ii) and 437∆ (panel iii) as inner membrane-tethered homo-tetramer, cytoplasmic homo-tetramer and cytoplasmic dimer, respectively. Large and small domains of each NTH are represented as buttons of different size connected by a linker in crossed conformation within a principal dimer, while the CTH of each protomer in the wild-type enzyme is shown as a wavy line emanating from the small domain and attached at its MTS motif to the inner membrane.

An allosteric regulatory site in RNase E-NTH is known as the 5’-sensor pocket (21, 22), with residue R169 located within it. This site confers to the enzyme the property of preferential action on RNA substrates with 5’-monophosphate termini (24, 25), which is lost with an R169Q substitution in the enzyme (26, 27). There is evidence for cross-subunit catalysis within a principal dimer, that is, with 5’-end sensing in one protomer and endonucleolytic cleavage in the adjacent one (21, 28). Newly synthesized bacterial RNAs bear 5’-triphosphate ends that need to be converted to 5’-monophosphate prior to RNase E action, and an enzyme RppH is required for this conversion (29, 30).

Notwithstanding that the C-terminal half of RNase E (CTH, residues 530 to 1061) is dispensable for viability, this region exhibits several important functional features and properties. The CTH is a scaffold for assembly of a multi-protein complex called the degradosome (31–37) whose constituents have been proposed to act cooperatively for efficient RNA degradation (38, 39). The CTH also bears two Arg-rich RNA-binding domains AR1 and AR2 (residues 604 to 644, and 796 to 814, respectively) (40, 41) as well as a membrane-targeting-sequence (MTS) motif from residues 565 to 582 (12, 42–46). The MTS motif is evolutionarily conserved in the gamma-and beta-proteobacteria (38, 42), and is responsible for the enzyme’s localization or compartmentalization to the cytoplasmic surface of the bacterial inner membrane. An F575E substitution within the MTS motif dissociates RNase E from the membrane and renders it soluble (20, 42, 43, 47). A recent study has also shown that the protein MinD binds to the MTS motif region of RNase E and by doing so mediates both the latter’s interaction with the membrane as well as the polar localization of a subset of mRNAs in *E. coli* (48).

That the CTH contributes significantly to RNase E function and hence to bacterial fitness and growth is supported by data from multiple studies [reviewed in (49)]. The specific activity of RNase E-ΔCTH is less than that of the full-length polypeptide both in vivo and in vitro (28, 50, 51), and mutants expressing the former are defective for degradation of ribosome-free mRNAs (52, 53). Loss of the MTS motif is by itself correlated with failure of degradosome localization to the inner membrane, reduced growth rate, and increased degradation of rRNAs (12, 42, 44). A transcriptomic study has shown that RNase E truncated beyond residue 602 (that is, with the MTS motif still present) is also defective for global mRNA degradation, which has been attributed to the loss of AR-1 and -2 domains in the mutant (54).

Importantly, perturbations of allosteric RNase E activation through its 5’- sensor pocket (caused either by absence of RppH or by an R169Q substitution in RNase E) are lethal in combination with RNase-ΔCTH but are viable with full length RNase E (27, 50). On the other hand, substitutions such as D303A or D346A that destroy the enzyme’s active site are lethal in both full-length and CTH-deleted RNase E (26, 28). These results have been interpreted to suggest that the enzyme catalyzes RNA cleavage through two pathways, namely a 5’-end-dependent pathway that requires R169 in RNase E and RppH, and a 5’-end-independent pathway that requires the CTH (50, 53, 55). The details of the postulated latter pathway are unclear.

In vitro studies on RNase E have been difficult because of the protein’s membrane association, insolubility, and tendency to aggregate (47); accordingly, the majority of studies have been performed with water-soluble RNase E-ΔCTH. The measured K*_cat_* values for RNA cleavage are rather low at around 10 min^-1^ or less (25, 26, 56–58), which may not adequately account for the enzyme’s essentiality and actions on its multifarious targets in vivo. The K*_cat_* value for (detergent-solubilized) full-length RNase E in a reconstituted degradosome is around 8-fold higher than that for RNaseE-ΔCTH (57). However, there are no published studies on the enzyme’s catalytic activity in vitro in its membrane-associated state.

We have recently demonstrated a new genetic phenomenon of "recessive resurrection" in RNase E, for which too the CTH is required (28). Recessive resurrection refers to the unexpected situation wherein an RNase E hetero-tetramer with just one wild-type subunit and three catalytically dead subunits (bearing D303A or D346A active site-substitutions) apparently exhibits nearly identical enzyme activity in vivo as that of a wild-type homo-tetramer. In other words, if four wild-type RNase E protomers were to be assembled in two alternative ways, either as a single homo-tetramer or as four hetero-tetramers each with three other catalytically inactive protomers, the latter (that is, the group of four hetero-tetramers) would exhibit approximately four-fold higher enzyme activity in vivo than would the former. This phenomenon is observed with the full-length polypeptides; on the other hand, in a tetramer comprised of CTH-truncated polypeptides, the presence of subunits with D303A or D346A active-site disruptions confers the expected, and opposite, effect of dominant negativity (that is, "subunit-poisoning") (28).

In the present study, we have performed studies of *rne-lac* expression and of RNA-Seq to confirm the phenomenon of recessive resurrection in RNase E, and to investigate which if any of the various features and properties of the CTH are required for its exhibition. Our findings suggest that it is the MTS motif that is mainly necessary for the purpose. Thus, the phenomenon of recessive resurrection is markedly impaired when the inactive RNase E protomers lack the MTS motif or bear the F575E substitution within the motif. Our model is that when RNase E is membrane-associated, it is rendered significantly more processive for its endonucleolytic action on RNA; and that recessive resurrection is simply a manifestation of processivity of catalysis by a homo-oligomeric enzyme.

## Materials and Methods

### Bacterial strains and growth conditions

Genotypes of *E. coli* K-12 strains are listed in Supplementary Table S1. The Δ*recA* allele was sourced from the Keio knockout collection (59). Cultures were routinely grown at 37° in LB medium (60). The medium was supplemented, where appropriate, with ampicillin (Amp), chloramphenicol (Cm), kanamycin (Kan), trimethoprim (Tp), and Xgal at concentrations as previously described (50). Supplementation with isopropyl β-D-thiogalactoside (IPTG) was at concentrations as indicated for the individual experiments, and it was ensured that all strains grown with IPTG were devoid of *lacY*-encoded lactose permease.

### Plasmids

Previously described plasmids included the following (salient features given in parentheses): pBR322 (Tet^R^ Amp^R^) (61); pKD13 (Kan^R^ Amp^R^), pKD46 (Amp^R^), and pCP20 (Cm^R^ Amp^R^) for use in recombineering and site-specific excision between FRT sites, as described (62); pSG1 and pSG4 (Kan^R^, pSC101-derived plasmids carrying, respectively, *rne^+^* and *rne*-R169Q alleles) (27); pHYD5159 and pHYD5193 (Tp^R^, single-copy-number *rne^+^* shelter plasmids with, respectively, *lacZ4526 lacY^+^* and *lacZ^+^* Δ*lacY* alleles) (28); pASKA-*rne* and its parent vector pCA24N from the ASKA plasmid collection (Cm^R^, wherein RNase E with N-terminal 6 X His-tag is expressed from an IPTG-inducible promoter in the former) (63); and pHYD5151, pHYD5152, pHYD5164 and pHYD5165 (Cm^R^, derivatives of pASKA-*rne* encoding RNase E mutants with, respectively, D346A, D303A, D346A + R169Q, and D303A + R169Q substitutions) (28).

Additional variants of pASKA-*rne* were obtained in this study by recombineering or by site-directed mutagenesis; they are listed in Supplementary Table S2 and their construction and validation are described in the Supplementary Text. The primers used for construction and validation of the plasmids are listed in Supplementary Table S3.

### Blue-white screening assay to determine functionality of *rne* alleles

The blue-white assay to establish whether an individual ASKA plasmid-borne *rne* allele can confer viability to a Δ*rne* strain has been described earlier (28). Briefly, a Δ*lac* Δ*rne* strain GJ20314 carrying the single-copy-number *rne^+^ lacZ^+^* Tp^R^ shelter plasmid pHYD5193 was transformed with the ASKA plasmid of interest and plated at suitable dilution on Xgal-supplemented medium without Tp. The presence of white colonies (typically at around 5% of total) indicated that cells that had spontaneously lost pHYD5193 continued to be viable and therefore that the ASKA *rne* allele was functional, whereas the absence of white colonies indicated that the *rne* allele was unable to support viability.

### Quantitation of *rne-lac* expression

β-Galactosidase assays were performed, with cultures of ASKA plasmid-bearing Δ*recA rne-lac* derivatives, by the method of Miller (60), and specific activity values were calculated in the units defined therein. For a given strain, *rne-lac* expression values at different IPTG concentrations were normalized to that at nil IPTG that had been determined in the same experiment, and data from at least three independent experiments were used for statistical analyses.

### Western blotting

The procedure for Western blotting was essentially as described (64). Cell lysates prepared from cultures grown to mid-exponential phase were subjected to electrophoresis on 8% polyacrylamide gels with sodium dodecyl sulphate. The separated proteins were electroblotted on PVDF membranes and reacted with mouse anti-His primary antibody followed by goat anti-mouse secondary antibody conjugated with horseradish peroxidase. The membranes were also stained with amido black to serve as loading controls.

### RNA-Seq experiments and data analysis

RNA-Seq experiments were performed with derivatives of strain GJ15074 [chromosomal genotype *rne^+^ rne-lacZ*(Δ*Y*)*A* Δ*recA*], each carrying a different ASKA plasmid expressing either wild-type RNase E or its variants with D303A, D346A, ΔMTS, or ΔMTS-D303A modifications. Cultures of the five strains were each grown in 200 ml of LB to early exponential phase; from each culture, 100 ml was transferred to a second pre-warmed flask already containing IPTG for a final concentration of 100 µM, and the pair of flasks (without and with IPTG) was then incubated for a further 30 min before being shifted to an ice bath. Each of the ten flasks was then spiked with 2 ml of a culture of GJ15074/pBR322 that had been freshly grown to mid-exponential phase in LB, after which cells from the cultures were harvested by centrifugation, washed once with single-strength TBS buffer (64), and stored at −80° until the time of their processing for RNA isolation. Two more batches of ten flasks each were similarly processed on succeeding days, so that triplicate RNA samples could be obtained for each strain without and with IPTG supplementation. [Of the total of fifteen pairs of cultures so prepared, one pair (replicate 3 of GJ15074/pASKA-*rne^+^*, without and with IPTG) was subsequently found to be unsatisfactory for its RNA yield and quality and was not further analyzed.]

The steps of RNA isolation from the frozen cell pellets and of RNA-Seq data generation were outsourced to Medgenome Labs Ltd (Bengaluru, India). As part of these steps, the RNA preparations were depleted of rRNA, and strand-specific RNA-Seq data (150 bases per read) were obtained from paired-end libraries on an Illumina sequencing platform.

Methods for RNA-Seq data analysis were largely as described earlier (65), with the following modifications. The Rockhopper algorithm (66) was used for alignment of reads to different genes of the MG1655 reference genome [(67), Accession number U00096.3] as well as to pBR322; the aggregate read counts aligned to MG1655 and to pBR322 for the different cultures are given in Supplementary Table S4. In a given culture, normalized read counts for each gene were determined as = [0.1 + (raw read counts aligned to the gene * 10^6^) / (gene length in bp * read counts for spiked-in pBR322 in the culture)].

Squared correlation coefficients (*R*^2^) were determined from pairwise comparisons of data from replicate cultures, and these values are given in Supplementary Table S5. For each of the strains and growth conditions (without or with IPTG), average normalized read counts were determined from the replicate pair with highest *R*^2^ value (values for all chosen pairs were ∆ 0.86). All rows were deleted for which the average normalized read values were < 3 in both IPTG-unsupplemented and –supplemented cultures of the strain with pASKA-*rne^+^*.

Alignment to successive 100-bp regions of the MG1655 genome was achieved with Bowtie-2 and SAM tools software (68, 69), and read counts were normalized to pBR322 read counts as described above. Heat maps were generated in R with the aid of the pheatmap package (70), using *k*-means clustering.

### Other methods

The protocols of Sambrook and Russell (64) were followed for recombinant DNA manipulations, PCR, transformations, and plasmid DNA analyses. Strain XL1-Blue (71) was routinely used for transformation of DNA preparations for new plasmid constructions. Procedures for phage P1 transduction (72), ARMS-PCR (73), recombineering (62), and pCP20-mediated site-specific excision between FRT sites (62), were as described.

## Results

### Exploiting *rne-lac* autoregulation to estimate intracellular RNase E specific activity

As mentioned above, the 5’-untranslated region of *rne* mRNA is itself a target for cleavage by RNase E, as a consequence of which the overall intracellular enzyme activity is subject to feedback autoregulation in *E. coli.* The expression of an *rne-lac* fusion therefore serves as a robust and sensitive inverse measure of intracellular RNase E activity, as established by work from several groups (13–19).

In the first part of the present study to investigate the roles of different CTH features on recessive resurrection, we adopted the following approach. The polypeptide encoded from the gene construct resident at the chromosomal *rne* locus (expressed from its native promoter) carried the functional active site residues D303 and D346, and hence was by itself sufficient to confer viability. Another gene construct placed downstream of an IPTG-inducible promoter on a plasmid encoded a second RNase E polypeptide that was either catalytically active (as control) or catalytically inactive (with D303A or D346A substitutions). Recessive resurrection was inferred to occur if there was an IPTG-dependent decrease of *rne-lac* expression in derivatives that carried the gene construct for a catalytically inactive polypeptide on the plasmid, since it implied that the catalytic activity of the functionally proficient polypeptide expressed from the chromosome is being augmented by its heteromeric association with functionally deficient polypeptides from the plasmid. On the other hand, if there was an IPTG-dependent increase of *rne-lac* expression under these conditions, the phenomenon was identified as one of dominant negativity.

### Description of the different RNase E constructs obtained and tested for recessive resurrection

ASKA plasmid-borne gene constructs for the following altered versions of RNase E (depicted in Fig. 1A) were obtained and validated as described in the Supplementary Text: truncations of the polypeptide beyond residues 815, 796, 645, 603, 529, or 437 [each designated by the corresponding residue number suffixed with Δ; 437Δ has been described as 438Δ in an earlier study (28)]; ΔAR1-ΔAR2 (that is, combination of individual deletions of AR1 and AR2); ΔMTS; ΔMTS -ΔAR1-ΔAR2; or the substitution F575E. [As schematically depicted in Figure 1B (panels i to iii), full-length RNase E and its truncated versions 529Δ and 437Δ are expected to exist as membrane-tethered homo-tetramer, cytoplasmic homo-tetramer, and cytoplasmic homo-dimer, respectively.] The different perturbations above were generated on constructs carrying the functional catalytic active site of RNase E or with D303A/D346A substitutions.

To investigate whether recessive resurrection would occur also in dimeric RNase E, we generated constructs to encode deletions of the small domain (since this domain mediates tetramer assembly as a dimer of principal dimers). In these ΔSD variants, the stretch of residues from 438 to 529 was deleted and replaced with an 8-amino acid sequence linking the large domain (and residues involved in Zn^2+^-coordination) to the CTH. The ΔSD constructs were generated on polypeptides carrying wild-type active site or the D303A substitution.

All constructs were tested by the blue-white screening assay described above to confirm that (i) those encoding the wild-type active site are able to confer viability in a chromosomal Δ*rne* strain background and (ii) those encoding D303A/D346A substitutions are unable to do so. Plasmid-bearing constructs encoding the functional active site were also introduced into a Δ*rne* strain (also RecA-deficient) carrying *rne-lac*, to obtain a measure of their intrinsic intracellular RNase E activity (Supp. Fig. S1). The data indicated that CTH truncations starting from residue 530 or further downstream (in each of which the enzyme is expected to remain tetrameric) are associated with moderate elevation of *rne-lac* expression (up to 4-fold), whereas the 437Δ truncation and the ΔSD construct (the latter was tested in a *recA^+^* derivative) conferred around 10-fold increase in *rne-lac* expression indicative of poor intrinsic enzyme activity. Perturbations to the MTS motif, as well as the ΔAR1-ΔAR2 deletion, were also associated with moderately elevated *rne-lac* expression.

### Testing for recessive resurrection with full-length and CTH-truncated RNase E polypeptides

In the studies described below, we have measured the values for autoregulated *rne-lac* expression within sets of isogenic *rne-lac* Δ*recA* strain derivatives carrying equivalent plasmid-based RNase E constructs, without or with a functional catalytic active site, expressed from an IPTG-regulated promoter; in the first set of experiments, all strains were *rne^+^* on the chromosome. For each construct, the *rne-lac* expression values at different IPTG concentrations have been normalized to that at nil IPTG (when the plasmid-encoded polypeptide is expected to be present at very low basal levels). It was also confirmed that *rne-lac* values at nil IPTG are similar for the various plasmid constructs (including vector pCA24N) in the given *rne^+^* chromosomal background (Supp. Fig. S2).

With the use of a trio of plasmid-borne full-length RNase E constructs (bearing either the functional active site or the D303A or D346A substitution therein), we first confirmed the phenomenon of recessive resurrection as previously described (28). Thus, in the chromosomal *rne^+^* background, the plasmid encoding functional full-length RNase E downstream of an IPTG-inducible promoter conferred, as expected, a progressive decrease in chromosomal *rne-lac* expression with increasing IPTG concentration, indicative of *rne* autoregulation; given that the extent of reduction in *rne-lac* expression was approximately 60 to 70 percent at IPTG concentrations ≥ 50 µM (Fig. 2, panel i), one may infer that the ratio of RNase E polypeptide synthesis from the plasmid to that from the chromosome is around 3:1 under these conditions. The relative excess of plasmid-encoded protomers in comparison to the chromosomally encoded species is expected to be accentuated also by the fact that the latter is expressed from the native autoregulated locus (and is therefore down-regulated to the same extent as *rne-lac*) whereas the former is induced by IPTG but not autoregulated. A nearly identical pattern of decrease in *rne-lac* expression was observed also with IPTG-induced expression of full-length RNase E variants with the catalytically dead D303A or D346A substitutions, indicative of recessive resurrection (Fig. 2, panel i). As expected, in the control strains carrying just the plasmid vector, *rne-lac* expression was unaffected by IPTG (Fig. 2, panel i).

**Figure 2:**
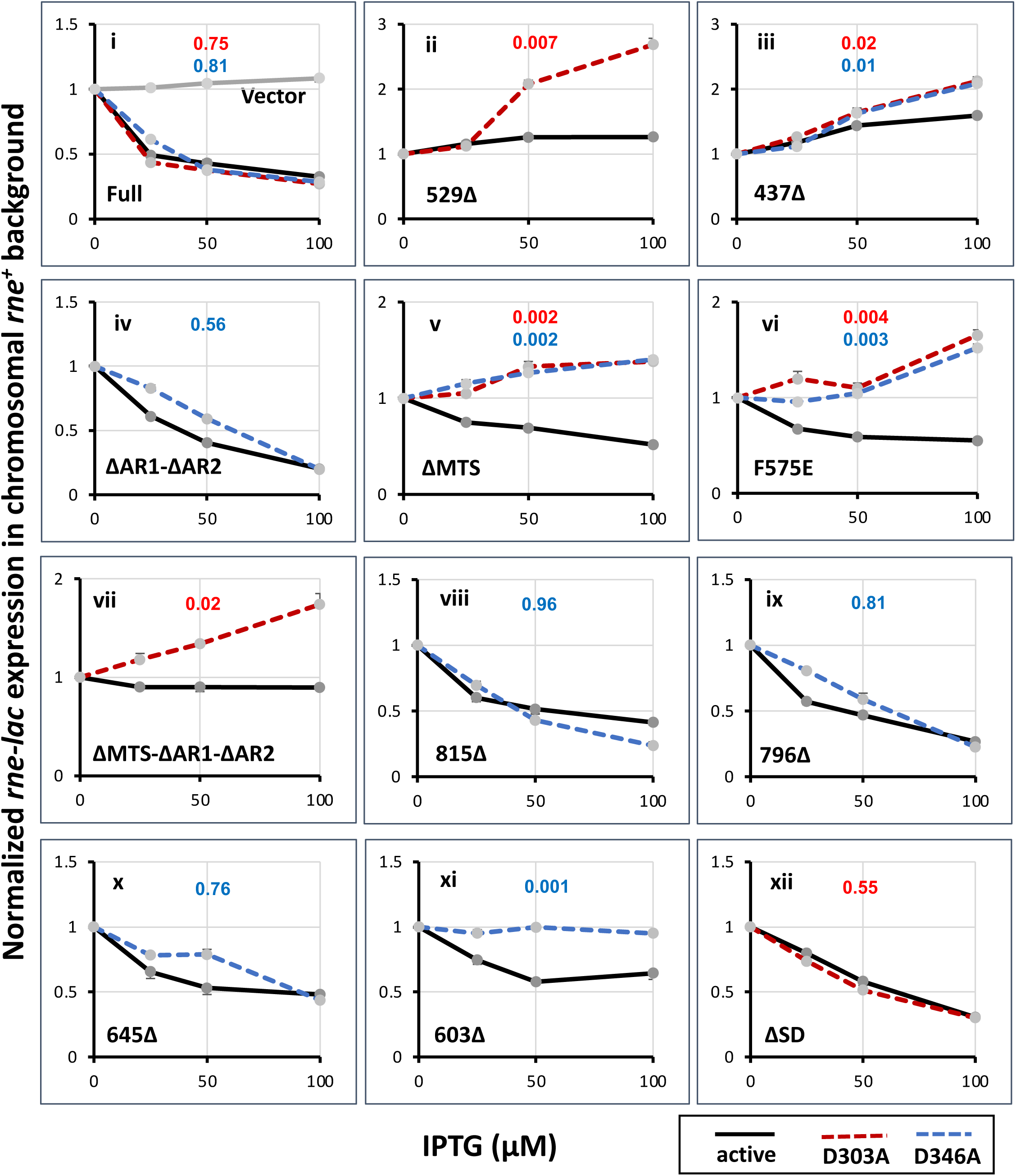
Effects on *rne-lac* expression in chromosomal ι1*recA rne^+^* background (strain GJ15074) of IPTG-induced overexpression of RNase E variants from ASKA plasmid constructs. In each panel, data are presented for a set of constructs (with functional active site or with D303A/D346A substitutions, as indicated in the key), and the identifying feature for each of the different sets is indicated at bottom left of the individual panels; Full, full-length. Values of β-galactosidase specific activity for a particular strain have been normalized, as explained in the Methods, to the value at nil IPTG for that strain (taken as 1.0), and standard error bars are included; the Miller unit values at nil IPTG for the different strains are given in Supplementary Figure S2. In each panel, *P* values are shown in red and blue, respectively, for the D303A and D346A variants, which were calculated as described in the text by Student’s *t*-test for the null hypothesis that the variants are not inferior to the cognate polypeptides bearing a functional active site. Also shown in panel i (grey line) are normalized values for the isogenic derivative bearing ASKA plasmid vector pCA24N.

In *rne-lac* expression experiments with plasmid-encoded RNase E polypeptides (with functional active site or with D303A or D346A substitutions) bearing 529Δ or 437Δ truncations (Fig. 2, panels ii and iii respectively), we confirmed the earlier findings that recessive resurrection is not observed with RNase E variants lacking the CTH (28). In either case, IPTG-induced expression from the plasmid of the control polypeptide with functional active site was associated with elevated *rne-lac* expression (more pronounced for 437Δ than for 529Δ), which is consistent with previous reports that C-terminal truncations of RNase E confer a dominant-negative phenotype (28, 51). Upon IPTG-induced expression of the polypeptides with D303A or D346A substitutions, the increase in *rne-lac* expression was even more marked, suggestive of subunit poisoning leading to stronger dominant negativity (which is to be viewed as the opposite of recessive resurrection). Indeed (and as further elaborated below), an important conclusion from multiple experiments in this study has been that recessive resurrection and dominant negativity are mutually exclusive manifestations associated with expression of different RNase E constructs bearing the D303A or D346A active site substitutions.

The qualitative interpretations above were supported also through statistical analyses, that are reported below for data obtained from cultures supplemented with 100 µM IPTG (see Fig. 2). The null hypothesis was that augmentation of intracellular RNase E activity (which is inversely correlated with *rne-lac* expression) upon overexpression of the polypeptides with D303A or D346A substitutions is not inferior to that upon overexpression of the cognate polypeptide with functional active site. The null hypothesis was rejected for the 529Δ and 437Δ constructs (*P* ≤ 0.01, Student’s one-tailed *t*-test), but not so for the full-length *rne* constructs (*P* = 0.30 and 0.21 for, respectively, D303A and D346A). For the full-length constructs, we also performed the converse statistical test of equivalence as described (74, 75; see also https://cscu.cornell.edu/wp-content/uploads/equiv.pdf); this analysis showed that intracellular RNase E activity upon overexpression of full-length polypeptides with D303A or D346A substitutions is not less than 60% of that upon similar overexpression of wild-type RNase E (*P* < 0.05 for both D303A and D346A).

### Recessive resurrection is impaired in the absence of an intact MTS motif in RNase E

The subsequent experiments of *rne-lac* expression were intended to test which features, if any, of the CTH are required for recessive resurrection. Our earlier studies had suggested that degradosome assembly on the CTH is dispensable for recessive resurrection (28); in continuation, the current experiments were performed with constructs bearing deletions of AR1 and AR2, or perturbations of MTS such as ΔMTS or F575E substitution. The results indicate that recessive resurrection is retained upon loss of the AR1 and AR2 motifs, in that IPTG-induced expression of the construct with D346A substitution and ΔAR1-ΔAR2 was associated with reduction in *rne-lac* expression in the *rne^+^* background (Fig. 2, panel iv). On the other hand, when the MTS motif was perturbed by either deletion or F575E alteration, constructs with the intact functional active sites were associated with IPTG-dependent reduction in *rne-lac* expression whereas those with D303A or D346A active site substitutions were dominant negative (Fig. 2, panels v and vi, respectively, for ΔMTS and F575E). These results suggest that, of the different features in the CTH, it is the MTS motif that is required for recessive resurrection.

Interestingly, with constructs bearing ΔAR1-ΔAR2 and ΔMTS together (that is, the triple deletion), IPTG-induced overexpression of even the polypeptide bearing the functional active site failed to lower *rne-lac* expression, and that with D303A conferred dominant negativity (Fig. 2, panel vii). These results suggest that the AR1-AR2 motifs and the MTS motif perhaps act redundantly to facilitate RNase E’s catalytic activity.

In a statistical analysis of the data obtained with 100 µM IPTG supplementation, the null hypothesis (that the increase in intracellular RNase E activity upon overexpression of the D303A-or D346A-bearing polypeptides is not lower than that obtained upon overexpression of the cognate construct with functional active site) was rejected for the constructs with ΔMTS, F575E, or ΔMTS-ΔAR1-ΔAR2 perturbations (*P* < 0.05, Student’s one-tailed *t*-test), but not so for the construct with ΔAR1-ΔAR2 (see Fig. 2). In the latter case, the statistical test of equivalence (74, 75) was employed to establish that the activity conferred by overexpression of this construct with D303A was not less than 60% of that obtained upon similar overexpression with the construct bearing the functional active site (*P* < 0.05).

### Comparable expression of RNase E polypeptides in strains without or with recessive resurrection

To exclude the possibility that the absence of recessive resurrection with some plasmid-encoded RNase E constructs is because of instability of the cognate RNase E polypeptides, we performed Western blot experiments following IPTG-induced expression from ASKA plasmid derivatives (with functional active site or with D303A / D346A substitutions) of full-length, 529Δ, ΔMTS and F575E versions of RNase E, in a chromosomal *rne^+^* background. The ASKA plasmid-encoded polypeptides are N-terminally His-tagged, and an anti-His antibody preparation was used in the experiments.

From the results (shown in Supp. Fig. S3), we confirmed that full-length RNase E (but not 529Δ) displays anomalously slow electrophoretic mobility, as reported earlier (76); the ΔMTS-and F575E-bearing polypeptides retained this property. It was also noted that the polypeptides that fail to show recessive resurrection, namely the D303A-and D346A-substituted versions of F575E (lanes 7-8) and of ΔMTS (lanes 10-11) are not less abundant than either (i) their counterparts with functional active sites (lanes 6 and 9, respectively) or (ii) the active site mutants of full-length RNase E which exhibit recessive resurrection (lanes 4-5). With the 529Δ constructs as well, expression of the polypeptide with D303A substitution (which is deficient for recessive resurrection) was similar to that with functional active site (compare lanes 12 and 13); interestingly, the abundance of both these polypeptides was markedly higher than those of the full-length, ΔMTS or F575E versions (after adjustment for amount of total protein loaded in each of the lanes).

We conclude that exhibition of recessive resurrection or otherwise is not correlated with differences in expression levels of polypeptides with D303A/D346A active site substitutions. This conclusion is supported also by data from the RNA-Seq experiments described below.

### Studying recessive resurrection through a global transcriptomic approach

The data from *rne-lac* expression studies described above had indicated that, in a strain bearing *rne^+^* at it native chromosomal locus, plasmid-directed overexpression of full-length RNase E polypeptides that are either catalytically active or catalytically dead lead to equivalent increases in intracellular RNase E activity, which we have interpreted as evidence for the phenomenon of recessive resurrection. Although the fact is well established that autoregulated *rne-lac* expression serves as a faithful reporter for the global activity in vivo of RNase E on its multiple individual substrates (13–19), the alternative possibility nevertheless had to be considered that our unexpected observations following overexpression of the catalytically inactive polypeptides were in some way specific to the *rne-lac* fusion transcript alone. Accordingly in the second part of this study, we adopted a global transcriptomic approach to examine the phenomenon of recessive resurrection, as outlined below.

The strains chosen for the RNA-Seq studies were the same as those that had been used for the *rne-lac* expression measurements. Thus, all of them were *rne^+^*, *rne-lacZ*(ι1*Y*)*A* and ι1*recA* on the chromosome, and each carried an ASKA plasmid derivative for IPTG-inducible overexpression of wild-type RNase E or one of its variants. As described in more detail in the Methods, cultures were grown in triplicate to early exponential phase in LB medium; one-half of each culture was then supplemented with IPTG, and cells from each of the cultures (without or with IPTG) were harvested 30 min later, together with a spiked-in aliquot of cells carrying the plasmid pBR322. RNA preparations from the cultures were depleted of rRNA and subjected to RNA-Seq (strand-specific, paired-end reads) on an Illumina platform. Thus, we sought in this approach to identify changes in transcript abundance caused by transient overexpression of RNase E or its variants in an *rne^+^* strain; it was assumed that the actions of PNPase and other 3’-5’ exoribonucleases (3, 4, 7, 9, 10, 77, 78), following endonucleolytic cleavage by overexpressed RNase E, would lead to a drop in transcript abundance in the IPTG-supplemented cultures.

RNA-Seq studies following transient perturbation of intracellular RNase E activity have previously been undertaken by several groups in *E. coli* and *S. enterica* (1, 77, 79) as well as in other Gram-negative bacteria [reviewed in (80)]. In each of these cases, a thermosensitive *rne* mutant was upshifted to the restrictive temperature to transiently inactivate RNase E, whereas in the present study we were performing the opposite kind of perturbation by transiently overexpressing RNase E or its variants in an *rne^+^* background. Thus, our study was expected to identify a different pattern of transcript abundance alterations caused by additional endonucleolytic cleavages than those that occur in the wild-type strain, at sites that are less efficiently recognized by RNase E. [RNA-Seq studies involving transient overexpression of toxin endoribonucleases have been reported earlier by Laub and coworkers (81, 82).]

Previous studies (1, 54, 77, 79, 83) have also shown that with reduced RNase E function, not only is there the expected increase in abundance of a large subset of RNAs which are the presumed substrates of the enzyme, there is also a decrease in abundance of a second subset. The numbers in these two subsets are comparable, but the aggregate magnitude of increased abundance is greater than that of the decrease. An unambiguous explanation for the latter subset (of decreased abundance) is lacking; it has been suggested that RNase E inactivation could have indirect effects through the stabilization of sRNAs or of mRNAs encoding transcription factors or other RNases (77, 83), or that certain RNA regions may be rendered stable following RNase E cleavage (77). Therefore, in the present study as well, we expected to observe both decrease and increase in abundance of different subsets of transcripts upon transient RNase E overexpression. In the context of recessive resurrection, the hypothesis under test was whether the pattern (and magnitude) of such alterations provoked with the wild-type enzyme would also be mimicked by overexpression of the catalytically dead D303A/D346A polypeptide variants.

### Support from RNA-Seq data analysis for the concept of recessive resurrection in RNase E

As described in the Methods, we first determined, from the RNA-Seq data for each strain and growth condition (without or with IPTG), average normalized read counts for the different protein-coding transcripts, sRNAs and tRNAs and the values (for 2971 genes, after removal of the gene rows with read counts below threshold) are given in Supplementary Table S6. The normalization was performed with respect both to gene length and to read counts aligned to the multicopy plasmid pBR322, which had been obtained following spike-in (at 2% v/v) of each of the cultures with a log-phase culture of the plasmid-bearing strain. With these data, we confirmed the following expectations from the strain genotypes: absence of reads for *recA* and *lacY*; approximate 100-fold increase in read counts for the *rne* coding region in IPTG-supplemented cultures; and, for the cultures heterozygous for *rne*, evidence for co-expression of *rne^+^* and the mutant *rne* allele, respectively, from the chromosome and from the ASKA plasmid.

We computed the ratio (designated as M) of the average normalized read count with IPTG to that without IPTG for each of the genes in a strain. M therefore represents the magnitude of change in RNA abundance for a gene upon transient overexpression of the ASKA-encoded polypeptide (with negative and positive log_2_ M values, respectively, for transcripts whose abundance decreases or increases following IPTG addition).

Analysis of the RNA-Seq data indicated that IPTG-induced overexpression of wild-type RNase E is associated with both negative and positive log_2_ M values for different genes, which is consistent with observations from the earlier studies that had employed mutants lacking RNase E (1, 77, 79, 83). Using a list of genes that exhibited log_2_ M >|1.73| for the strain with ASKA-*rne_+_*, (that is, an absolute change in RNA abundance of >3-fold in cultures with IPTG relative to that without IPTG), we prepared two heat maps of log_2_ M values (across all five strains) for the genes whose transcript abundance was, respectively, decreased (Fig. 3A upper image, 174 genes) or increased (Fig. 3A lower image, 121 genes) in that list; the two lists are presented in sheets 1 and 2, respectively, of Supplementary Table S7, wherein all values of log_2_ M >|1.73| in the different columns are marked in blue. (The *rne* gene itself was excluded from consideration in these analyses, since its greatly elevated log_2_ M value is simply a reflection of IPTG-induced overexpression of the gene from the ASKA plasmid.)

**Figure 3:**
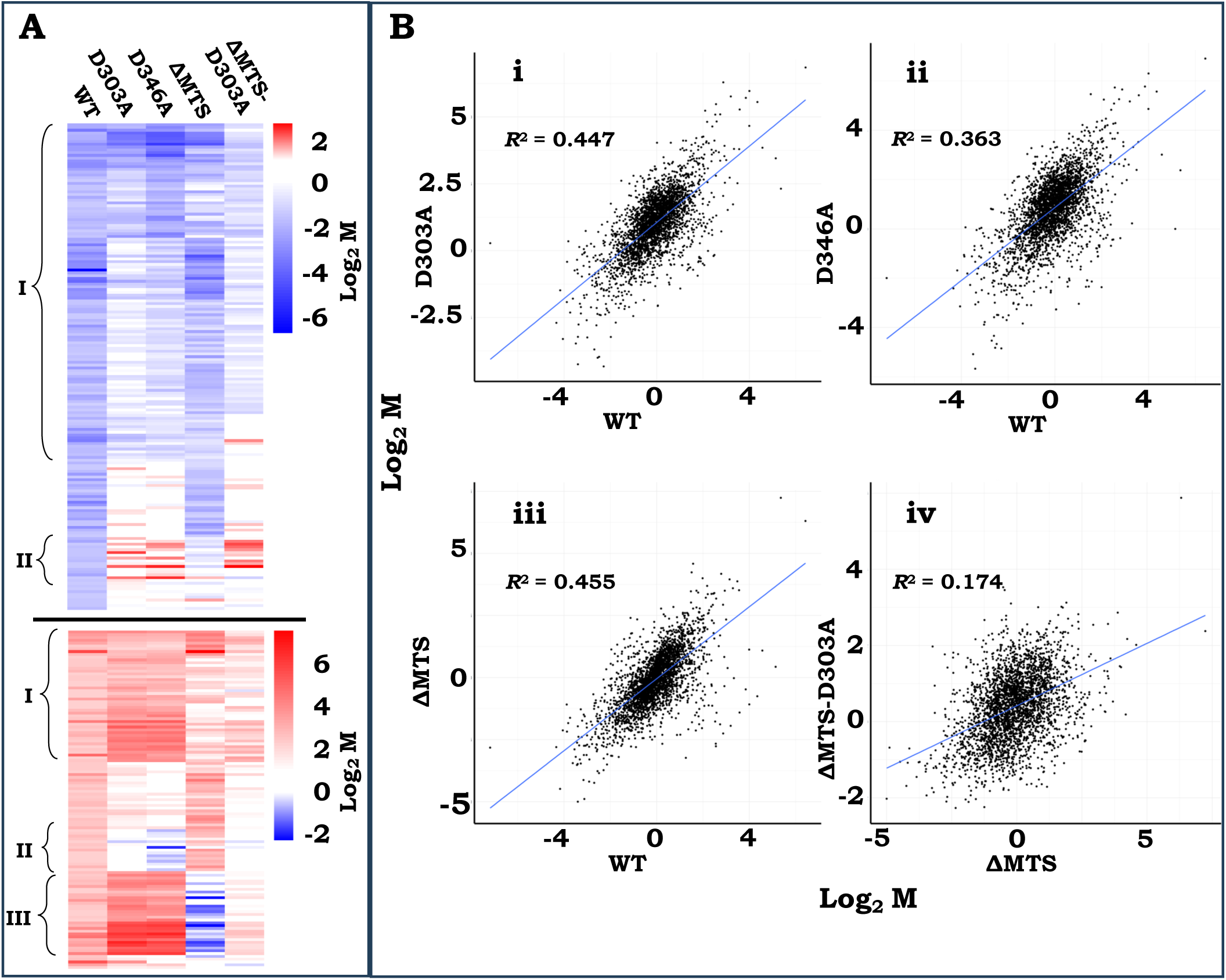
Analysis of RNA-Seq data from strains overexpressing wild-type (WT) RNase E or one of its variants (D303A, D346A, ∆MTS, and ∆MTS-D303A). (**A**) Heat map representations of log_2_ M values in the different strains for genes with log_2_ M values (in the strain with WT RNase E) < −1.73 or > 1.73 (upper and lower images, respectively, each with its own colour code as shown). Regions marked ’I’, ’II’, and ’III’ are explained in the text. (**B**) Log_2_ M-M plots are shown for the following strain pairs: i, WT vs. D303A; ii, WT vs. D346A; iii, WT vs. ∆MTS; and iv, ∆MTS vs. ∆MTS-D303A.

In both heat maps, a comparison between the columns for D303A and D346A on the one hand with that for wild-type RNase E on the other indicated the existence of two subsets of genes, that are demarcated as I and II in Figure 3A. The larger subset I corresponds to transcripts whose change in abundance was more or less concordant across the three strains (indicative of recessive resurrection), whereas the transcripts of subset II were discordantly affected. The two subsets are further discussed below and in the subsequent sections.

The log_2_ M-M plots, depicting the IPTG-dependent alterations in transcript abundance for all genes in the strain with ASKA-*rne^+^* on the one hand against those in the strains with ASKA-*rne*-D303A or ASKA-*rne*-D346A on the other, are presented in Figure 3B, panels i and ii, respectively. The effects of overexpression either of wild-type RNase E or of the mutant polypeptides (with D303A or D346A substitutions) were reasonably well correlated at the whole-genome level, with *R*^2^ values of 0.45 and 0.36, respectively. Notably, there were approximately 140 genes that concordantly exhibited log_2_ M >|1.73| in both (i) the cultures overexpressing wild-type RNase E as well as (ii) the cultures of at least one of the mutant-overexpressing strains (D303A or D346A). Of these, more than 100 were commonly affected in all the three strains (see Supp. Table S7), and some of them are further described below. The overlap of the downregulated genes in strains overexpressing the D303A or D346A variants compared to those in the strain overexpressing wild-type RNase E was statistically highly significant (*P* < 10^−35^ and < 10^−46^, respectively, hypergeometric test). The upregulated genes were also highly correlated between these strains, as described in a subsequent section.

Taken together, therefore, these results provide strong support for the concept, originated from the *rne-lac* studies, for the phenomenon of recessive resurrection in RNase E.

### RNA-Seq analysis in strains overexpressing RNase E-ΔMTS or RNase E-ΔMTS-D303A

The effects of overexpression of RNase E-ΔMTS and of RNase E-ΔMTS-D303A, each in the *rne^+^* chromosomal background, were also examined in the RNA-Seq experiments. Since, upon IPTG supplementation, the plasmid-encoded transcripts are in 100-fold excess over the transcripts expressed from chromosomal *rne^+^*, it is expected that the predominant enzyme oligomers of interest in the former case would be the homo-tetramers of overexpressed ΔMTS, whereas in the latter case they would be hetero-tetramers of overexpressed ΔMTS-D303A assembled with wild-type RNase E protomers in 3:1 ratio (since the homo-tetramers of ΔMTS-D303A themselves are catalytically dead). [Given that the determinants of RNase E tetramerization reside in the NTH of the polypeptide (21, 22), we have assumed that the ΔMTS protomers are equivalent to wild-type RNase E protomers for assembly into tetramers.]

In each of the two heat map representations [for genes exhibiting, respectively, decreased (Fig. 3A, upper image) and increased (Fig. 3A, lower image) transcript abundance upon transient overexpression of wild-type RNase E], the pattern for the ΔMTS variant was more or less aligned to that for wild-type RNase E with the exception of a subset in the latter (designated III in Fig. 3A) that was prominently discordant. As explained below, this subset corresponds to genes of the σ^32^-heat shock regulon.

We also prepared log_2_ M-M plots to compare the effects of overexpression (in the *rne^+^* chromosomal background) of RNase E-ΔMTS against wild-type RNase E on the one hand, and of RNase E-ΔMTS-D303A against RNase E-ΔMTS on the other (Fig. 3B, panels iii-iv, respectively). The correlation for the first comparison was reasonable (*R*^2^ = 0.46), and the overlap (with the strain overexpressing wild-type RNase E) of downregulated genes with log_2_ M < −1.73 was statistically highly significant (*P* < 10^−82^, hypergeometric test).

On the other hand, there was minimal correlation in the second comparison between the effects of overexpression of ΔMTS and of ΔMTS-D303A (*R*^2^ = 0.17); for the latter strain, there were only 17 genes with log_2_ M < −1.73, indicating that overexpression of ΔMTS-D303A had little consequence for the transcript-degrading activity associated with chromosomal *rne^+^*. These findings would suggest that the endonucleolytic activity in a hetero-assembly of ΔMTS-D303A : wild-type is simply proportionate to the number of wild-type protomers in the tetramer.

Thus, the foregoing analysis of RNA-Seq data permits the inferences (i) that recessive resurrection is manifested in the catalytic activity of RNase E across numerous substrates, and (ii) that this property is markedly impaired in the ΔMTS variant of the enzyme.

### Examples of multi-gene operons whose transcript abundance is concordantly reduced upon overexpression of wild-type RNase E or its D303A/D346A variants

The list in Table S7 includes several operons (such as *tdcABCDEF*, *srlAEBDM*, *garPLR*, *tnaAB*, and *treBC*) wherein the transcript abundance of multiple genes was concordantly diminished upon overexpression of either RNase E or its D303A/D346A variants. Log_2_ M plots for these operons at 100-bp resolution served to confirm these findings, and furthermore to show that the concordance largely extended to the situation of RNase E-ΔMTS overexpression but not to that of RNase E-ΔMTS-D303A overexpression (Fig. 4A-B and Supp. Fig. S4A-C). These data therefore serve to reaffirm, at the level of individual genes and operons, the conclusions drawn from *rne-lac* expression studies that recessive resurrection is exhibited in full-length RNase E but is impaired upon loss of the MTS motif in the polypeptide.

**Figure 4:**
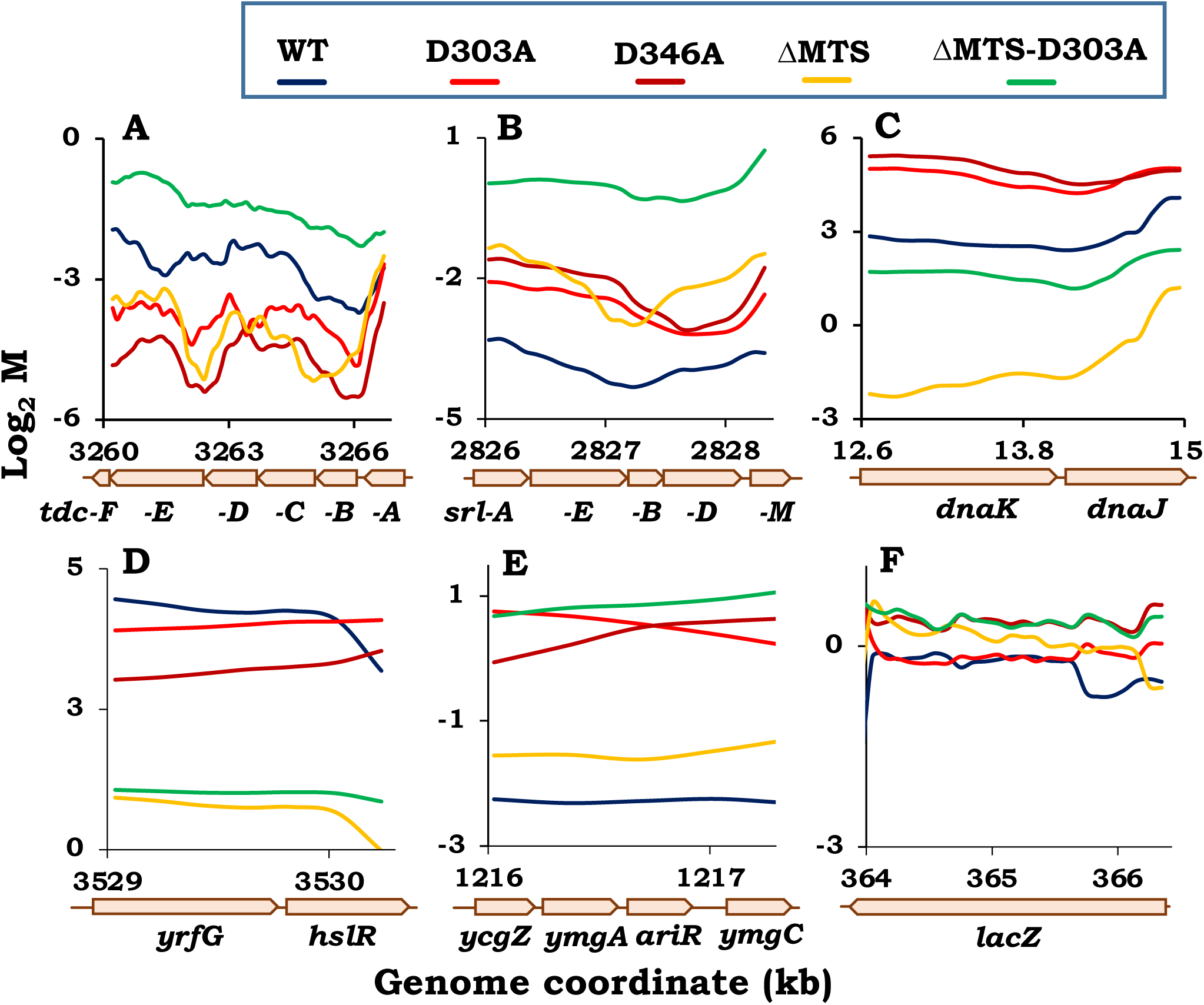
Plots of moving averages of log_2_ M values from RNA-Seq data at 100-base resolution for different genes in strains overexpressing wild-type (WT) RNase E or one of its variants (D303A, D346A, ∆MTS, and ∆MTS-D303A). Moving averages in each case were determined with a window of 9 and step size of 1. Arrows indicate gene extent and direction of transcription, and genome coordinates (in kb) are marked. Representative examples are shown of: (**A-B**) recessive resurrection in operons with negative log_2_ M; (**C-D**) genes of the σ^32^-heat shock regulon with positive log_2_ M; and (**E**) an operon exhibiting a discordant response upon overexpression of WT RNase E or its D303A/D346A variants. (**F**) Plot for the *lacZ* gene.

### Increased abundance of RNAs encoding heat shock proteins upon overexpression of RNase E or its D303A/D346A variants

The heat shock response and its regulation in *E. coli* have been subjects of extensive study earlier [reviewed in (84–86)]. A major player in heat shock response regulation is the RNA polymerase sigma factor σ^32^ or RpoH, whose overexpression is associated with transcriptional induction of 127 genes (87).

In our RNA-Seq experiments, of 74 genes whose transcript abundance was elevated > 3-fold (log_2_ M > 1.73) commonly upon RNase E overexpression as well as upon overexpression of both its inactive variants D303A and D346A, 41 (shown in red in Table S7, including *dnaKJ*, *groES-groEL*, *yrfG-hslR*, *hslVU*, and *ibpAB*) were common to the list of 127 genes induced by RpoH overexpression (87). The overlap between the two lists was statistically highly significant (*P* < 10^− 32^, hypergeometric test), and log_2_ M plots at 100-bp resolution are shown for some of them (Fig. 4C-D, and Supp. Fig. S4D). Most interestingly, transcript abundance of the heat shock genes was not elevated upon overexpression of the the ΔMTS variant of RNase E, and indeed for many of them (such as *htpG*, *clpB*, *ibpAB*, *dnaK*, *groES-groEL*, *hslRO*, and *hslVU*) the log_2_ M values were <−1 indicative of their downregulation (Table S7). The possible interpretations from these observations are discussed below.

### Examples of transcripts whose abundance is altered only upon overexpression of RNase E polypeptides with a functional active site

From the heat map representations, it was inferred that a subset of genes (demarcated II in Fig. 3A-B) were discordant for alterations in their transcript abundance between cultures with overexpression of wild-type RNase E or its ΔMTS variant on the one hand and those with overexpression of the active site mutants (D303A, D346A, or ΔMTS-D303A) on the other. This was also borne out by analysis of the log_2_ M data for the five strains. Listed in sheets 1 and 2 of Supplementary Table S8 are 86 genes (60 and 26 with reduced and increased abundance, respectively, upon wild-type RNase E overexpression and which are subsets of genes listed in sheets 1 and 2, respectively, of Table S7) that exhibited discordant behaviour between cultures with wild-type RNase E overexpression and those with overexpression of D303A or D346A variants [log_2_ M < −1.73 in former and > −0.3 in one or both of the latter; or > 1.73 in former and < 0.3 in one or both of the latter). For a majority of these genes, the effects of ΔMTS overexpression and of ΔMTS-D303A were similar, respectively, to that of wild-type RNase E and of its D303A/D346A variants.

As examples from the category of the discordant “reduced abundance” genes, log_2_ M plots at 100-bp resolution are depicted for the *ycgZ-ymgA-ariR-ymgC* operon (Fig. 4E) as well as for *katE* and *talA-tktB* (Supp. Fig. S4E-F). Furthermore, 17 of the 27 genes in the discordant “increased abundance” list were found to encode different tRNAs (shown in red in Table S8); their over-representation in this group (given that the *E. coli* genome contains 86 tRNA genes) was statistically highly significant (*P* < 10^−21^, hypergeometric test). The possible interpretations from these findings are discussed below.

### Absence of change in (*rne-*)*lacZ* RNA abundance upon transient RNase E overexpression

As mentioned above, the RNA-Seq experiments with transient IPTG-induced overexpression of RNase E (or its variants) were performed on derivatives carrying the *rne-lacZ*(Δ*Y*)*A* fusion on the chromosome. The specific activity of β-galactosidase in the *rne-lac* strain with ASKA-*rne^+^* was significantly reduced, as expected, in cultures supplemented transiently with IPTG (for 30 min) in comparison to the cultures without IPTG (Supp. Fig. S5), indicative of RNase E autoregulation. On the other hand, it was evident from the RNA-Seq data (Table S6) that *lacZ* mRNA abundance at the whole-gene level was not concomitantly reduced in the IPTG-supplemented cultures, although *lacA* transcript abundance did exhibit a reduction under these conditions.

The log_2_ M plots at 100-bp resolution for the *lacZ* gene in the different strains are presented in Figure 4F. In all strains, transcript abundance was unchanged across most of the length of *lacZ* following IPTG addition, which appears to be discordant with the β-galactosidase specific activity measurements. The findings may perhaps be explained by invoking the distinction between chemical decay and functional decay of transcripts (88–91), upon their cleavage by RNase E. The results with *rne-lacZ* also suggest that the catalog (in Table S7) of RNAs exhibiting reduced abundance (at the gene level) may be an underestimate of the actual numbers which may have been functionally inactivated upon transient RNase E overexpression.

### Effects on recessive resurrection of other truncations within the CTH

Given that our analysis of the transcriptome data above (with respect to full-length and ΔMTS constructs of RNase E) had validated the conclusions from the *rne-lac* expression experiments, we proceeded to employ the latter to test for recessive resurrection with other constructs and in other strain backgrounds. The results from these studies are described in this and the following sections.

As shown above (Fig. 2, panels ii-iii) and earlier (28), constructs of the RNase E truncations 529Δ and 437Δ (both lacking the MTS motif) do not exhibit recessive resurrection in the *rne-lac* expression assay, and we now asked whether less extensive truncations within the CTH such as 815Δ, 796Δ, 645Δ or 603Δ will do so. Our findings indicate that as with full-length RNase E, the 815Δ, 796Δ and 645Δ constructs (all of which retain the MTS motif) are able to confer recessive resurrection in the *rne^+^* background (Fig. 2, panels viii-x, respectively), and this was also established by the statistical test of equivalence that was performed as described above (with data from 100 µM supplementation, *P* < 0.05 in all cases).

On the other hand, with 603Δ, *rne-lac* expression was reduced upon IPTG addition only for the construct with functional active site but not for that with D346A substitution, suggestive of absence of recessive resurrection in this situation (Fig. 2, panel xi); the null hypothesis that the two polypeptides confer similar effects upon overexpression with 100 µM IPTG was rejected (*P* = 0.03, Student’s one-tailed *t*-test). Since the length of C-terminal extension beyond the MTS motif in the 603Δ polypeptide is only 21 amino acid residues, it is likely that its propensity for membrane association is compromised.

Taken together, therefore, the results of *rne-lac* expression with the different CTH truncations are also consistent with the proposal that it is the MTS motif in RNase E that is required for the phenomenon of recessive resurrection.

### Recessive resurrection is exhibited in dimeric RNase E

The quaternary structure of the NTH of RNase E is that of a dimer of principal dimers, and it is the pair of small domains (between residues 438 and 529) in a principal dimer that contacts its counterparts of the second principal dimer for homo-tetramer assembly. To determine whether recessive resurrection can occur in dimeric RNase E that is still membrane-associated, we constructed and employed the plasmid-borne, and IPTG-regulated, ΔSD variants (Δ438-529), in each of which the SD region is deleted and the CTH region is attached directly to the large domain (along with the segment involved in Zn^2+^ co-ordination) through an 8-amino acid Gly-rich linker (92). When a ΔSD variant is overexpressed in a chromosomal *rne^+^* strain, the full-length polypeptide is expected to be titrated by the ΔSD species so that the former is present predominantly in hetero-dimeric assemblies that are membrane-associated.

As mentioned above, in a Δ*rne* background, the ΔSD variant (with functional active site) was associated with high *rne-lac* expression (Supp. Fig. S1), indicating that it possesses low specific activity (although sufficient to support viability). It is not clear, however, whether this low activity is because of the protein’s existence as a dimer (rather than as tetramer) or because of alteration of its primary structure through presence of the artificial linker region between the large domain and CTH.

In experiments with a pair of plasmid-borne ΔSD variants (carrying the functional active site or the D303A substitution) in the chromosomal *rne^+^* background, we could show that *rne-lac* expression is decreased with increasing IPTG to equivalent extents with both the variant bearing the functional active site as also that in which the active site had been destroyed (Fig. 2, panel xii). The answer to a statistical test for equivalence (74, 75) was in the affirmative, that intracellular RNase E activity associated with the ΔSD-D303A variant upon 100 µM IPTG supplementation is not less than 60% of the activity obtained with the ΔSD construct carrying the functional active site (*P* < 0.05). These findings suggest that recessive resurrection (namely, the ability of a catalytically inactive protomer to contribute to enzyme activity in heteromeric assembly with a functional protomer) is exhibited even in dimeric RNase E.

### 5’-Monophosphate sensing is not required for recessive resurrection in RNase E

The R169Q substitution in RNase E which abolishes sensing of, and stimulation of catalytic activity by, 5’-monophosphate in the RNA substrate is non-lethal when present on the full-length polypeptide but is lethal in combination with a CTH truncation. Since recessive resurrection is also exhibited with full-length but not CTH-truncated RNase E, we investigated whether this phenomenon would still occur in heteromeric complexes in which all protomers carried R169Q.

Accordingly, *rne-lac* expression was measured in different derivatives of a strain in which full-length RNase E with R169Q substitution was expressed from its native promoter. The derivatives each carried a plasmid expressing one of the following from an IPTG-inducible promoter: (i) nil (that is, plasmid vector); (ii) full-length RNase E with functional active site or with D303A or D346A substitutions; or (iii) full-length RNase E with R169Q substitution along with functional active site or with D303A or D346A substitutions.

From the results, we noted first that the basal *rne-lac* expression (that is, in presence of just the plasmid vector) is elevated around three-fold in the derivative with RNase E-R169Q compared to that with wild-type RNase E (Supp. Fig. S6, compare the first two bars at left). This finding lends support to the notion that the 5’-sensing and 5’-independent pathways contribute redundantly to catalytic efficiency of the enzyme in vivo (26, 50).

With regard to recessive resurrection, we observed that in the derivatives with RNase E-R169Q, the pattern of reduction in *rne-lac* expression upon IPTG-induced expression of different constructs is very similar across all combinations of the alterations tested (that is, with or without functional active site and with or without 5’-sensor mutation), and especially so at 50µM IPTG (Fig. 5). This interpretation was supported also by statistical tests of equivalence, that each of the variants conferred not less than 60% of the activity conferred by wild-type RNase E upon their overexpression with 50 µM IPTG (*P* < 0.05 in all cases). These data suggest that recessive resurrection is exhibited even when all four protomers are defective for 5’-monophosphate sensing.

**Figure 5:**
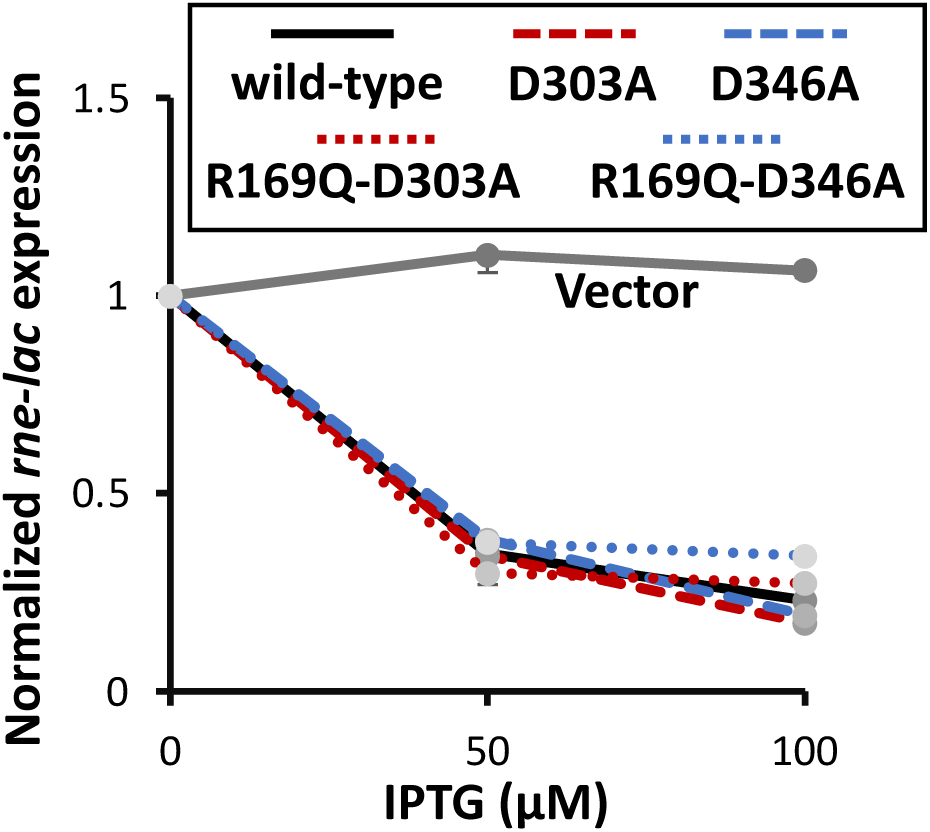
Effects on *rne-lac* expression in ∆*recA rne*-R169Q background (strain GJ20323/pSG4) of IPTG-induced overexpression of RNase E variants from ASKA plasmid constructs. Notations are as described in legend to Figure 2. Miller unit values at nil IPTG for the different strains are given in Supplementary Figure S6.

## Discussion

Genetic, biochemical, and structural studies have provided reasonably detailed insights into the mechanism of catalysis by the NTH of the essential endoribonuclease RNase E of *E. coli*. The role of its CTH in the cellular functions of the enzyme is less well understood. We had previously identified a novel genetic feature of recessive resurrection in RNase E which requires the CTH of the polypeptide (28). Recessive resurrection refers to the phenomenon by which four wild-type RNase E protomers, each distributed in a hetero-tetramer with three other catalytically dead protomers, together apparently confer roughly four-fold higher enzyme activity in vivo than when assembled as a single homo-tetramer. The present work suggests that it is the MTS motif in the CTH that is needed for recessive resurrection.

### Identification of the MTS motif as a requirement for recessive resurrection in RNase E

In this work, we have sought to compare intracellular enzyme specific activity of a catalytically active RNase E homo-tetramer with that of a hetero-tetramer in which one, two, or three protomers carried the D303A or D346A substitution inactivating the catalytic active site. Such hetero-tetrameric assemblies were achieved by progressive increase in expression of the inactive polypeptide from an IPTG-inducible promoter on a plasmid, whereas the wild-type protomer bearing the functional active site was expressed from the *rne* promoter at its native chromosomal locus. Intracellular RNase E activity was assessed through two approaches: (i) by measurement of steady-state *rne-lac* expression (which is inversely related to RNase E activity, since the latter is autoregulated); and (ii) by determination of changes in abundance of individual transcripts following transient overexpression of different RNase E polypeptides in a chromosomal *rne^+^* strain.

The results from both approaches have indicated that recessive resurrection is exhibited in full-length RNase E, and that the MTS motif in the polypeptide is required for the purpose. In other words, loss of a functional active site (by either D303A or D346A substitution) confers different effects depending upon whether it has occurred in full-length RNase E or in its ΔMTS version: in a chromosomal *rne^+^* strain, overexpression of the mutant polypeptide in the former case is associated with elevation of intracellular RNase E activity to roughly the same extent as that obtained with wild-type RNase E overexpression; whereas in the latter case, there is no such elevation (and instead, there may even be a dominant negative effect).

Furthermore, with the approach employing *rne-lac* expression, it was shown that a single substitution (F575E) which disrupts the MTS motif is sufficient to impair recessive resurrection. There was a correlation between presence of the MTS motif in different constructs and their ability to support recessive resurrection (the sole exception being the 603Δ construct, whose deficiency for recessive resurrection may perhaps be explained by the MTS motif’s location close to the C-terminal end of the polypeptide). Finally, the phenomenon was also exhibited in dimeric RNase E bearing the CTH, although less prominently than in the tetrameric constructs.

### How is membrane anchoring of RNase E mediated by the MTS motif?

Disruption of the evolutionarily conserved MTS motif in RNase E (by either deletion or F575E substitution) has previously been shown to be associated with moderate reductions in both mRNA degradation rate and culture growth rate (12, 42). It has also been proposed that membrane localization of RNase E and of the degradosome serves both to protect against degradation of precursor rRNAs within ribosome assembly intermediates in the cytoplasm (44), and to mediate the capture of polyribosomes for promoting mRNA degradation (31). How would these findings and proposals align with the new results, that recessive resurrection in RNase E hetero-tetramers with only one functional catalytic active site requires presence of the MTS motif in the protomers?

From the X-ray crystal structures of the NTH of RNase E, it is apparent that the junction between NTH and CTH is located on the inner lateral surface of the small domain, that is, adjoining its partner small domain in each of the principal dimers of the tetramer. This would imply that the MTS motifs from the four protomers of each RNase E molecule are sterically clustered during their interaction with the bacterial inner membrane (as depicted in Fig. 1B, panel i), given that the MTS motif is only around 35 amino acid residues away from the NTH-CTH junction in the primary sequence of the polypeptide. This model is different from the extended-degradosome-conformation models depicted previously (23, 49), but it may also explain the results from live-cell microscopy experiments in which RNase E molecules were visualized to form short-lived foci or puncta at the membrane (2, 31, 43).

It may be expected that in the ΔMTS mutant, both this clustering and the membrane association are disrupted so that the individual CTH regions of a tetramer are more widely separated from one another in a three-dimensional random-walk distribution. Accordingly, we propose that it is the clustered arrangement of the CTH regions of all four protomers mediated by the MTS motifs that is required for the phenomenon of recessive resurrection in RNase E (Fig. 1B, panel i), as further discussed below.

### Postulated mechanism for recessive resurrection, and its contrast with dominant negativity

From the results depicted in Figure 2, it would appear that when different versions of catalytically dead RNase E protomers are overexpressed in cells with a basal level of functional full-length protomers, the phenomenon of either recessive resurrection or dominant negativity is elicited in a mutually exclusive fashion. These two phenomena are characterized by reduction and elevation, respectively, of *rne-lac* expression with progressive overexpression of the catalytically inactive protomers in the cells. Thus, dominant negativity was observed with inactive versions of CTH-truncated polypeptides such as 529Δ or of polypeptides with ΔMTS or F575E alterations, whereas recessive resurrection was exhibited with inactive versions of full-length, 645Δ, or ΔAR1-ΔAR2 polypeptides.

We propose that the contrasting phenomena of recessive resurrection and of dominant negativity can be explained by, respectively, (i) processive degradation of RNA at the single-molecule level by membrane-associated RNase E, and (ii) distributive or stochastic cleavage of RNA at single sites by the cytoplasmic enzyme. Below, we discuss first the distinction between processive versus distributive activities (93) with respect to RNA cleavage by RNase E, and then the postulated effects of the enzyme’s subcellular localization on these alternative events.

It has long been speculated (94) that endonucleolytic cleavages in *E. coli* occur progressively in a 5’ to 3’ direction on mRNA, and that such decay is often cotranscriptional (14, 95). Evidence in strong support of these notions was obtained from an RNA-seq study by Xie and coworkers (96) on lifetimes of different segments of transcripts across the genome. Belasco and coworkers (97, 98) have also shown that the endonucleolytic action of RNase E in vivo likely involves a linear 5’ to 3’ scanning mechanism of the enzyme on RNA. However, it is not known whether the polarity of RNA degradation is being achieved by a processive cleavage mechanism, or by a successive series of distributive cleavages on the RNA molecules (24).

According to our model, endonucleolytic activity of a wild-type RNase E tetramer within the cell is processive in 5’ to 3’ direction on a single molecule of RNA substrate, as may be represented by the following scheme:

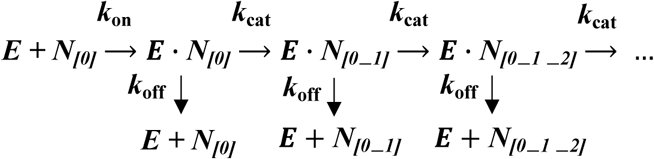

In the scheme shown above, (i) *E* and *N_[0]_* stand for the *e*nzyme and initial length of the *n*ucleotide substrate, respectively; (ii) *N_[0_1]_*, *N_[0_1_2]_*, and so on represent the successively smaller lengths of substrate after, respectively, the first, second steps and so on of the processive RNase E-catalyzed cleavage reactions (the released oligoribonucleotides in these reactions are not shown); and (iii) *k*_on_, *k*_off_, and *k*_cat_ have the usual meanings that specify the rates of enzyme-substrate association, dissociation, and catalysis, respectively.

Our proposed scheme for processive action of RNase E is analogous to that described earlier for other enzymes such as hydrolases of cellulose and chitin (99–101) or nucleic acid enzymes including helicases (102), nucleases (103) and polymerases (104, 105); the kinetics of catalysis by these enzymes are distinct from the classical Michaelis-Menten kinetics. Thus, for such enzymes and as is also proposed here for RNase E, there is a component of substrate retention and channeling so that (i) the rate-limiting step of the reaction is determined not by the enzyme’s *k*_cat_ but often by its rate of dissociation *k*_off_ from the substrate (99–101, 104, 105), and (ii) the enzyme’s *k*_cat_ merely determines its processivity, which can be expressed as the ratio of *k*_cat_ to *k*_off_ (99, 101).

By our model, therefore, although the presence of three inactive protomers in a tetramer of full-length RNase E will be expected to decrease the enzyme’s *k*_cat_, such reduction would serve to reduce enzyme processivity but not the rate of RNA decay. That the CTH of RNase E may participate in substrate trapping and presentation for catalysis has been suggested earlier (28, 51, 57), but our proposal that the CTH-bearing enzyme acts processively on the substrate is novel. Recessive resurrection is, therefore, a phenomenon that is an apparent outcome of the enzyme’s postulated processivity on RNA in vivo.

We further propose that the enzyme variants which display dominant negativity are those that exhibit high *k*_off_ rates; in this situation, processivity is lost and the classical Michaelis-Menten kinetics become applicable, with *k*_cat_ becoming rate-limiting. Accordingly, a reduction in *k*_cat_ caused by presence of inactive protomers within a tetramer will lead to drastic decrease in rate of RNA degradation, accounting for dominant negativity. In summary, *k*_off_ and *k*_cat_, respectively, are proposed to be rate-limiting for membrane-associated RNase E that exhibits recessive resurrection and cytoplasmic RNase E that exhibits dominant negativity.

In our model, how could membrane association of RNase E be expected to contribute to its processivity? As indicated above, we believe that the MTS motif mediates the clustered arrangement of the CTH regions of a tetramer in the vicinity of the membrane. The *E. coli* inner membrane is comprised of acidic and zwitterionic phospholipids (106, 107), and it has previously been shown that such a membrane is strongly repulsive for nucleic acid molecules such as DNA (108). It is therefore possible that it is electrostatic repulsion between RNA and the inner membrane (both negatively charged) that leads to a reduced rate of dissociation of RNA from the enzyme-substrate complex, thereby contributing to the enzyme’s processivity.

Subcellular compartmentalization of endoribonucleases involved in mRNA degradation has been reported also in other bacteria (2), with the examples including membrane-anchored RNase Y in *Bacillus subtilis* and *Staphylococcus aureus* (109), and presence of RNase E in liquid-liquid phase-separated condensates in *Caulobacter crescentus* (110). Whether such compartmentalization may be correlated with enzyme processivity on mRNA remains to be investigated. Also worthy of investigation is the question whether other homo-oligomeric enzymes that display processivity, such as several of the exonucleases (103, 111) or helicases (112, 113), could exhibit recessive resurrection.

### Recessive resurrection in RNase E does not require RNA 5’-monophosphate sensing

It has previously been shown that RNA 5’-monophosphate sensing by RNase E is dispensable for viability when the polypeptide carries the CTH. This finding has led to the hypothesis of a second and redundant CTH-mediated pathway of presentation of RNA to the enzyme’s catalytic active site for cleavage. Our present results indicate that recessive resurrection (which is CTH-dependent) also does not require RNA 5’-monophosphate sensing, since it is exhibited even when all four polypeptides in a tetramer carry the R169Q substitution. The simplest explanation for these observations taken together is that it is the 5’-end-independent (and CTH-dependent) pathway which mediates the endonucleolytic processivity of RNase E on its substrate.

### Additional features associated with overexpression of RNase E and its variants

From the RNA-Seq data and their analyses, two major additional findings were (i) the identification of transcripts whose abundance is altered upon overexpression only of RNase E polypeptides with functional active site (that is, wild-type or ΔMTS) but not of polypeptides bearing active site substitutions, and (ii) the demonstration that transcripts of the σ^32^-heat shock regulon are upregulated upon overexpression of the full-length polypeptides (wild-type or its D303A/D346A variants) but not of the ΔMTS versions. The transcript categories so implicated correspond to subsets II and III, respectively, in the heat map representations (Fig. 3A).

The former category may be viewed as comprising genes that are not subject to recessive resurrection. We suggest that in these cases, efficient endonucleolytic cleavage of the concerned substrates might require the co-operative actions of more than one catalytically active protomer within an RNase E tetramer. With regard to tRNAs that exhibited discordant changes in abundance, one possibility is that overexpression of RNase E polypeptides with functional active site (but not of those with D303A/D346A substitutions) leads to an enhancement of processing of the cognate primary or precursor tRNA transcripts (8, 10, 114) as to result in their increased abundance.

To account for the latter category of upregulated heat shock genes, we suggest that overexpression of RNase E is associated with enhanced degradation of a particular mRNA or sRNA that then results in increased levels of RpoH. The finding that upregulation of these genes occurs even upon overexpression of the D303A/D346A variants is because of recessive resurrection. It is also necessary to assume that this particular mRNA or sRNA belongs to the category of substrates that are cleaved only by membrane-anchored but not cytoplasmic (that is, the ΔMTS variant of) RNase E (46), which feature might also in some way be tied to the fact that RpoH itself is associated with the inner membrane (85, 86).

An alternative possibility is that the generalized increase in endonucleolytic cleavage of mRNAs upon RNase E overexpression leads to increased synthesis of truncated misfolded proteins and consequently to induction of the heat shock response. However, we believe that this model is less likely, since it fails to explain why transcript abundance of the heat shock genes is not elevated upon ΔMTS overexpression, even though the effects of overexpression of wild-type RNase E and ΔMTS polypeptides are themselves well correlated.

## Data availability

All RNA-Seq data described in this work are available for full public access at https://www.ncbi.nlm.nih.gov/bioproject/PRJNA1169523.

## Supplementary data

Supplementary data are provided in a PDF file “Supplementary data”, along with three Excel files as Tables S6, S7 and S8, respectively.

## Acknowledgements

We thank Nida Ali and members of the Laboratory of Bacterial Genetics, Centre for DNA Fingerprinting and Diagnostics, Hyderabad for strains; Ayushi for assistance with isolation of and studies on the ΔSD mutant; and Rachna Chaba, Purnananda Guptasarma, Himanshi Maheshwari and Nalini Raghunathan for advice and discussions. This paper is dedicated to the memory of Marc Dreyfus.

## Supplementary Text

### Construction of ASKA-*rne* plasmid variants

ASKA plasmids encoding different RNase E variants were constructed by recombineering or by site-directed mutagenesis, as described below. These variants were generated on ASKA-*rne* as well as its D303A / D346A-encoding derivatives pHYD5152 and pHYD5151, respectively. The oligonucleotide primers used are listed in Supplementary Table S3.

#### I. Construction by recombineering of ASKA-*rne* plasmid variants encoding C-terminal truncations of RNase E

Forward primers for preparing the variants encoding truncations 603Δ, 645Δ, 796Δ, and 815Δ were, respectively, PB18, PB19, PB20 and PB21; each of these primers carried a TAA stop-codon sequence immediately following the sequence for the cognate truncation site and preceding that for annealing to the recombineering plasmid pKD13. Primer G, described in a previous study (1), was used as the common reverse primer for preparing all the truncation variants. Primers J and S, that were used as forward primers for truncations 437Δ and 529Δ, respectively, have also been described earlier (1).

For generating each of the truncations, the pair of forward and reverse primers was first used in PCR on plasmid pKD13 (2), and the amplicon was then transfected into strain MC4100 carrying both plasmid pKD46 as well as plasmid ASKA-*rne* or its D303A / D346A-encoding variants. Kan^R^ colonies were selected, and derivatives carrying the truncation-encoding variant on the ASKA replicon were validated as described below.

#### II. Construction of ASKA-*rne* plasmid variants by site-directed mutagenesis

The procedure for site-directed mutagenesis to introduce mutations encoding in-frame internal deletions as well as the F575E substitution in the ASKA plasmid derivatives (ASKA-*rne*/pHYD5151/pHYD5152) was essentially as described (1). Twenty cycles of "linear PCR" were performed with Q5 DNA polymerase and the indicated primer pairs on the plasmid templates, and the amplicons were transformed following *DpnI* treatment into strain XL1-Blue with selection for Cm^R^ colonies.

The primer pairs employed for the different mutations were as follows: (i) ΔMTS, ME01-ME02; (ii) ΔAR1, ME07-ME08; (iii) ΔAR2, ME09-ME10; (iv) F575E substitution, ME11-ME12; and (v) ΔSD, ST1-ST2. The primers for generating the ΔSD construct are so designed that *KpnI* digestion of the amplicon and its re-circularization by ligation will yield a plasmid encoding an RNase E variant with deletion of the small domain and its replacement by a short Gly-rich linker peptide.

Combinations of deletions, such as ΔAR1-ΔAR2 and ΔMTS-ΔAR1-ΔAR2 were achieved in sequential steps with the primer pairs listed above.

### Tests for validation of new plasmid constructs

The tests performed to validate the new plasmid constructs are described separately below for those obtained by recombineering and those by site-directed mutagenesis. Two additional tests were commonly applied to the plasmids obtained through either approach: (i) inviability of MC4100 derivatives carrying the plasmids on medium supplemented with 100 µM IPTG, which served to exclude the occurrence of inadvertent frameshift mutations; and (ii) blue-white screening assay to confirm that plasmids with functional active site and those with D303A/D346A substitutions confer, respectively, viability and inviability in a Δ*rne* background.

#### (I). Plasmids encoding RNase E truncations obtained by recombineering

Since it was possible that the Kan^R^ recombineering event to generate truncations of RNase E could occur on either the ASKA plasmid or the chromosome, we first prepared plasmids from the strains and transformed them into XL1-Blue to check for co-inheritance of Cm^R^ and Kan^R^ markers. Agarose gel electrophoresis of undigested plasmid preparations from the XL1-Blue transformants was undertaken to exclude plasmid multimers, and presence of the truncation of interest was established both by restriction digestion with *BamHI* and by PCR [using a pair of primers of which one (K1) was common for validating all truncations and has been described in Datsenko and Wanner (2) while the other was either GJRNE04 (for 815Δ, 796Δ, 645Δ and 603Δ) or PB13 (for 529Δ and 437Δ)].

ARMS-PCR tests (3) were use to determine whether the residue at position 575 of RNase E was Phe (parent) or Glu (mutant) with primer pairs PB23-PB25 for the former and PB24-PB25 for the latter.

#### II. Plasmids obtained following site-directed mutagenesis

Presence of deletions ΔSD, ΔAR1, ΔAR2, and ΔMTS (either alone or in different combinations) was checked by (i) restriction digestion with *BamHI*; (ii) PCR with primer pairs ST3-ST4 for the ΔSD constructs, PB13-GJRNE08 for constructs with ΔMTS and/or ΔAR2 but with intact AR1, and ME05-GJRNE09 for constructs bearing ΔAR1 either alone or in combination with ΔMTS and/or ΔAR2; and (iii) DNA sequencing of the PCR amplicons above.

DNA sequence analysis of the two ΔSD constructs obtained in this study (with functional active site and with D303A substitution) revealed that, in both of them, the peptide linker replacing the small domain was eight residues in length (Gly-Thr-Ser-Gly-Gly-Gly-Gly-Ser), although the linker length that had been expected from the designed primer sequence was thirteen amino acid residues (Gly-Thr-Ser-Gly-Gly-Gly-Gly-Ser-Gly-Gly-Gly-Gly-Ser).

### ARMS-PCR testing for variants encoding R169Q, D303A or D346A substitutions

The primer pairs used for ARMS-PCR (3) to determine whether the residues at positions 169, 303 and 346 were parental or mutant were the following: position 169, PB34-PB37 and PB27-PB35 for parent and PB34-PB38 and PB27-PB36 for mutant; position 303, PB30-ST2 for parent and PB31-ST2 for mutant; and position 346, PB32-ST2 for parent and PB33-ST2 for mutant.

---

**Table S1.**
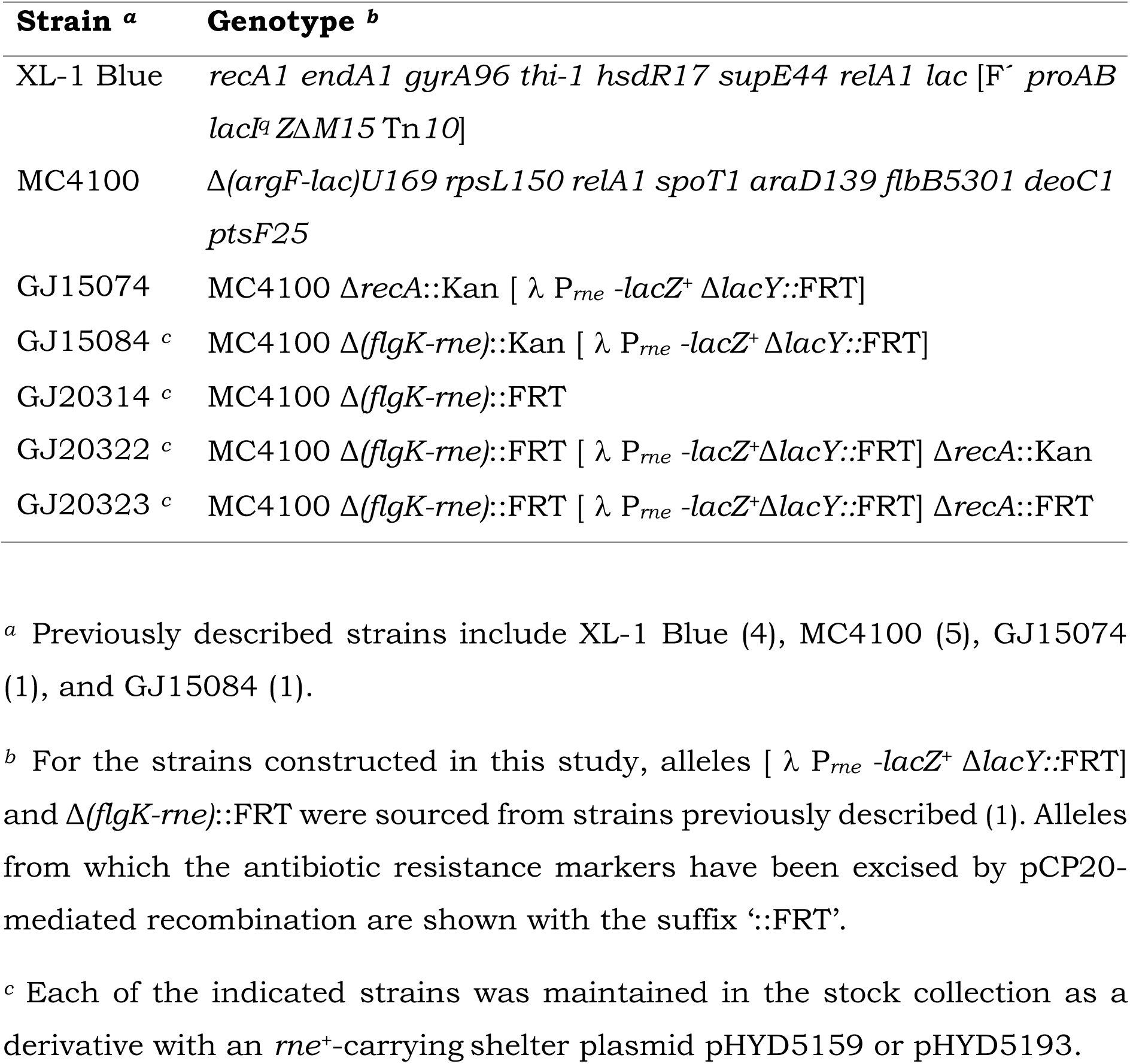
List of *E. coli* strains.

**Table S2.** Features of RNase E-encoding variants on ASKA plasmids that were constructed in this study.

| **Plasmid** | **Feature at enzyme active site** | **Additional feature(s)** |
| --- | --- | --- |
| pHYD7693 | Functional | ΔSD |
| pHYD7695 | D303A | ΔSD |
| pHYD7716 | Functional | ΔAR1, ΔAR2 |
| pHYD7717 | D303A | ΔMTS |
| pHYD7718 | D346A | ΔMTS |
| pHYD7719 | Functional | 603Δ::Kan |
| pHYD7720 | Functional | 645Δ::Kan |
| pHYD7721 | Functional | 796Δ::Kan |
| pHYD7722 | Functional | 815Δ::Kan |
| pHYD7723 | D346A | 603Δ::Kan |
| pHYD7724 | D346A | 645Δ::Kan |
| pHYD7725 | D346A | 796Δ::Kan |
| pHYD7726 | D346A | 815Δ::Kan |
| pHYD7732 | D346A | ΔAR1, ΔAR2 |
| pHYD7733 | D303A | 529Δ::Kan |
| pHYD7737 | Functional | 437Δ::Kan |
| pHYD7738 | D303A | 437Δ::Kan |
| pHYD7739 | D346A | 437Δ::Kan |
| pHYD7744 | Functional | F575E |
| pHYD7745 | D303A | F575E |
| pHYD7746 | D346A | F575E |
| pHYD7747 | Functional | 529Δ::Kan |
| pHYD7801 | Functional | ΔMTS |
| pHYD7804 | Functional | ΔAR1, ΔAR2, ΔMTS |
| pHYD7805 | D303A | ΔAR1, ΔAR2, ΔMTS |

**Table S3.**
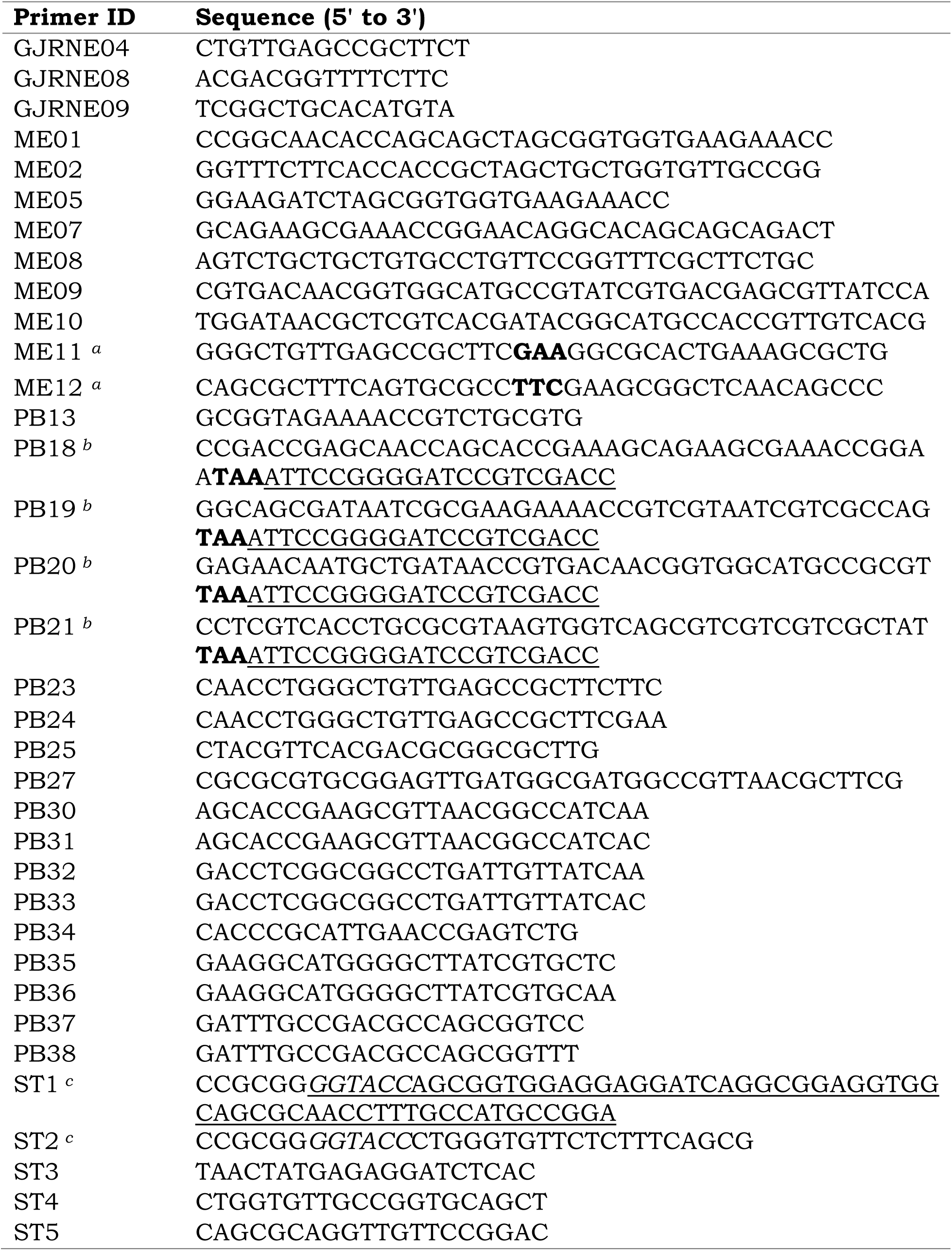
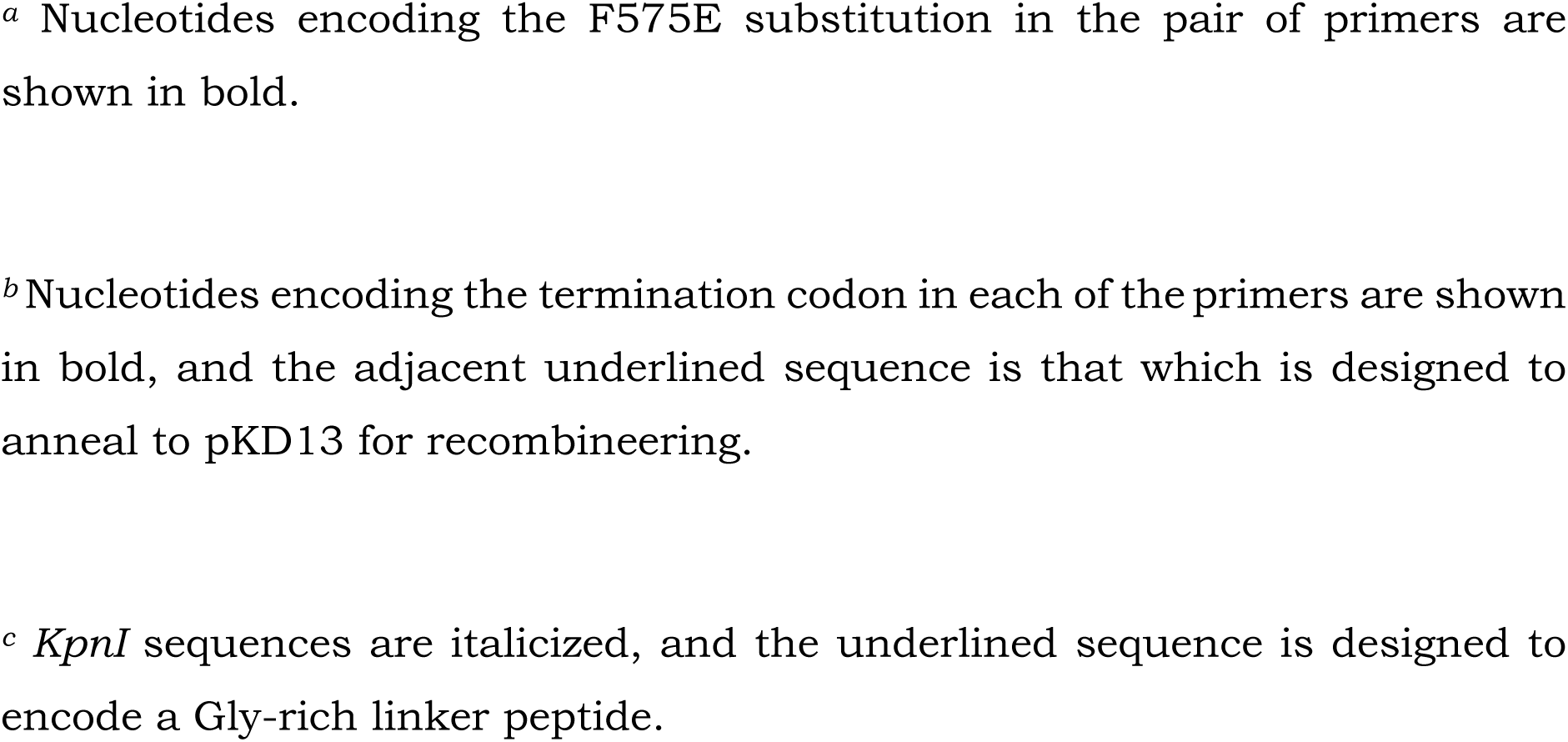
List of oligonucleotide primers.

**Table S4.** Aggregate RNA-Seq read counts aligned to chromosome (and aligned to pBR322 within parentheses) for different cultures.

| RNase E polypeptide encoded by ASKA plasmid | IPTG | Replicate culture |  |  |
| --- | --- | --- | --- | --- |
|  |  | 1 | 2 | 3 |
| wild-type | – | 4.2e6 (14379) | 4.8e6 (17984) | not available |
|  | + | 4.8e6 (13438) | 4.1e6 (12701) | not available |
| D303A | – | 4.5e6 (16387) | 3.3e6 (19315) | 7.6e6 (22090) |
|  | + | 3.8e6 (12419) | 6.4e6 (9127) | 7.8e6 (15370) |
| D346A | – | 6.8e6 (12586) | 7.0e6 (16542) | 5.8e6 (12317) |
|  | + | 6.8e6 (6934) | 6.5e6 (9486) | 8.0e6 (9409) |
| ΔMTS | – | 7.5e6 (17310) | 7.2e6 (12131) | 4.2e6 (10353) |
|  | + | 6.3e6 (10702) | 7.5e6 (14111) | 7.1e6 (9630) |
| ΔMTS-D303A | – | 5.4e6 (12361) | 6.6e6 (10233) | 6.4e6 (14138) |
|  | + | 7.1e6 (11030) | 6.4e6 (11490) | 10.1e6 (8952) |

**Table S5.** Pair-wise *R*^2^ values for RNA-Seq data from triplicate cultures (pairs used for the determination of averages in Table S6 are underlined)

| RNase E polypeptide encoded by ASKA plasmid | IPTG | Replicate pair |  |  |
| --- | --- | --- | --- | --- |
|  |  | 1-2 | 2-3 | 1-3 |
| wild-type | – | <u>0.95</u> | not available | not available |
|  | + | <u>0.92</u> | not available | not available |
| D303A | – | 0.61 | 0.57 | <u>0.89</u> |
|  | + | <u>0.93</u> | 0.66 | 0.63 |
| D346A | – | 0.94 | 0.94 | <u>0.95</u> |
|  | + | 0.81 | <u>0.86</u> | 0.62 |
| ΔMTS | – | <u>0.99</u> | 0.96 | 0.97 |
|  | + | 0.42 | <u>0.92</u> | 0.56 |
| ΔMTS-D303A | – | 0.93 | 0.90 | <u>0.99</u> |
|  | + | <u>0.91</u> | 0.63 | 0.83 |

**Supp. Fig. S1:**
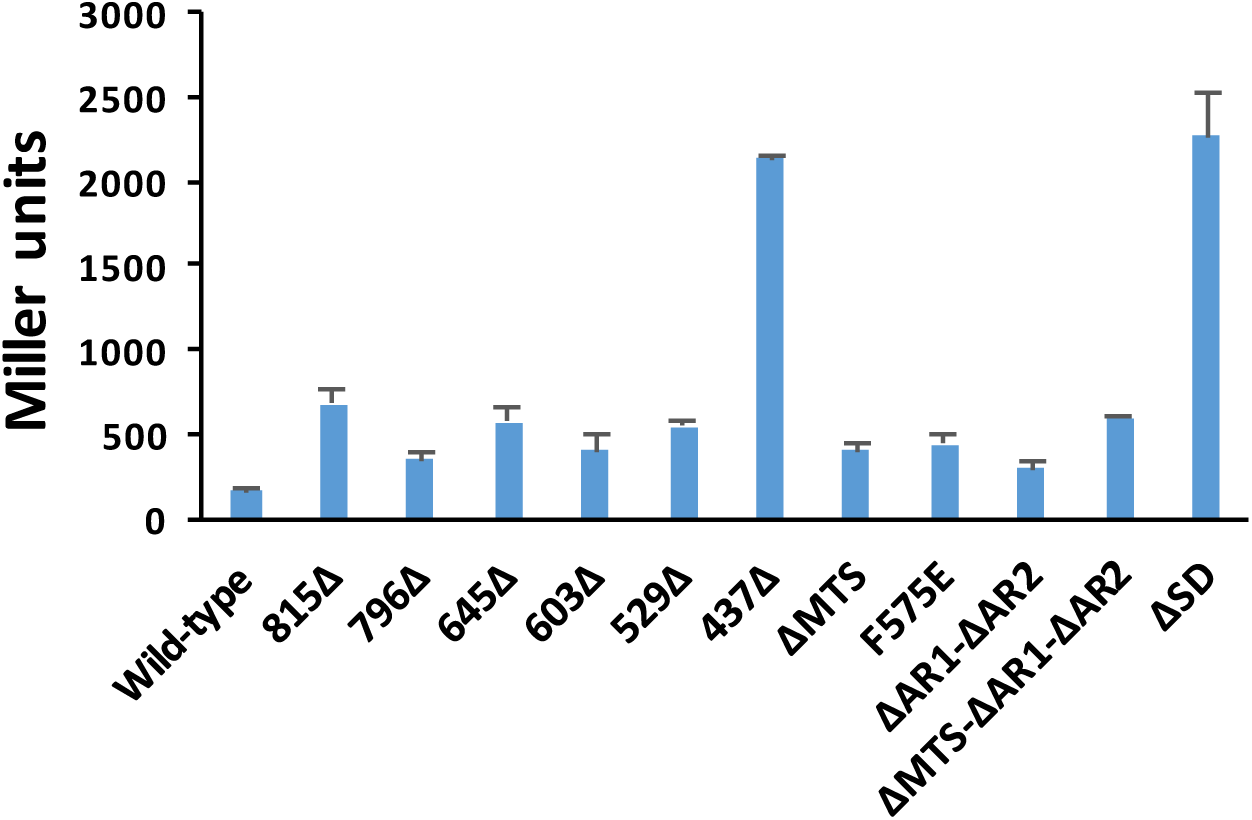
Specific activity (Miller units) of β-galactosidase in cultures supplemented with 20 μM IPTG of derivatives of Δ*rne rne-lac* Δ*recA* strain GJ20322 carrying different plasmids as indicated. The ΔSD-encoding plasmid variant was assayed in the Δ*rne rne-lac recA^+^* strain GJ15084. Plasmids were (from left, numbers are to be prefixed with pHYD unless otherwise marked): ASKA-*rne*, 7722, 7721, 7720, 7719, 7147, 7737, 7801, 7744, 7716, 7804 and 7693.

**Supp. Fig. S2:**
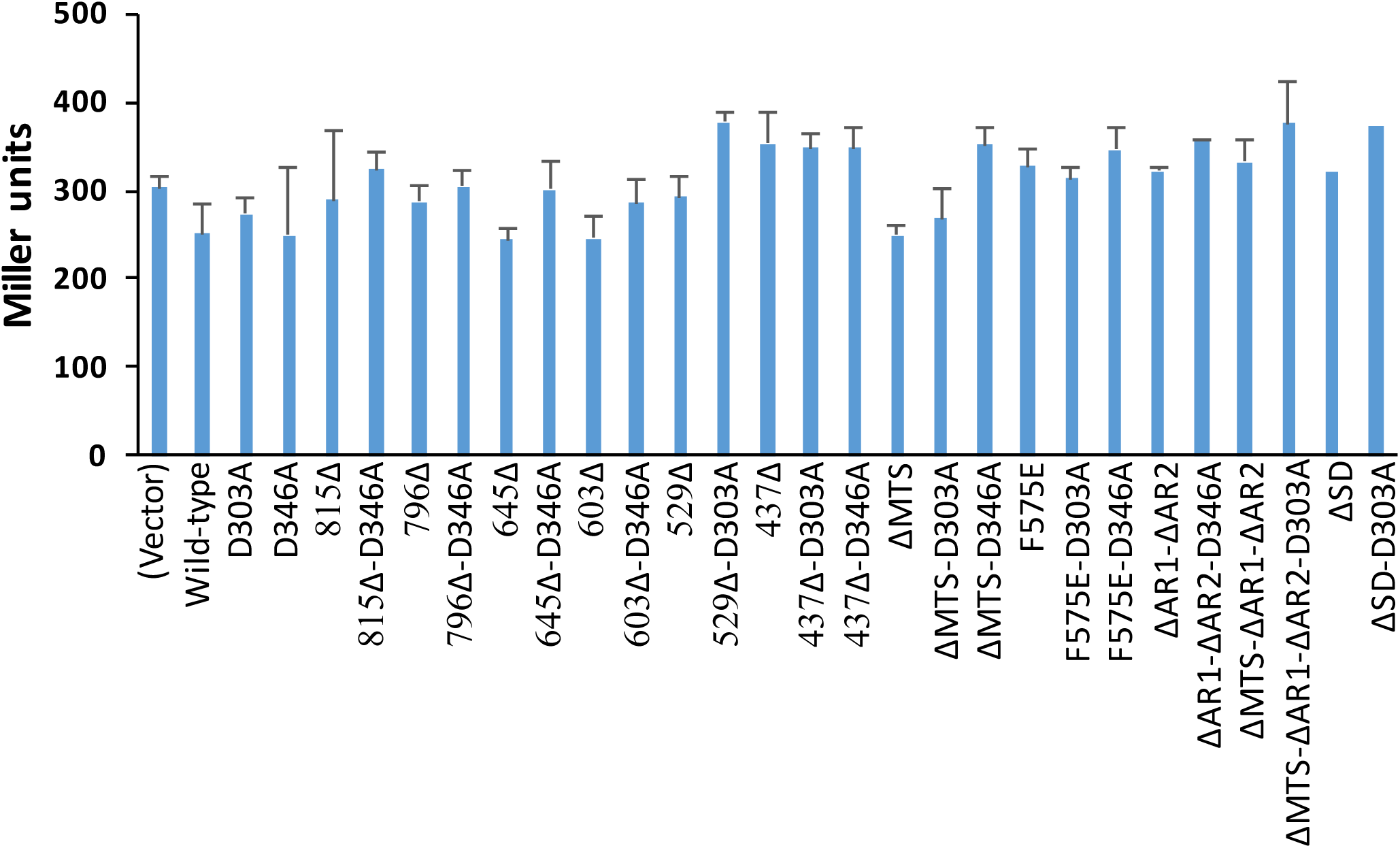
Specific activity (Miller units) of β-galactosidase in cultures supplemented with nil IPTG of derivatives of *rne^+^ rne-lac* Δ*recA* strain GJ15074 carrying ASKA plasmid vector or its derivatives enco ding different RNase E polypeptides as indicated. These values have been normalized to 1.0 in the different panels of Figure 2. Plasmids were (from left, numbers are to be prefixed with pHYD unless otherwise marked): pCA24N, ASKA-*rne*, 5152, 5151, 7722, 7726, 7721, 7725, 7720, 7724, 7719, 7723, 7747, 7733, 7737, 7738, 7739, 7801, 7717, 7718, 7744, 7745, 7746, 7716, 7732, 7804, 7805, 7693 and 7695.

**Supp. Fig. S3:**
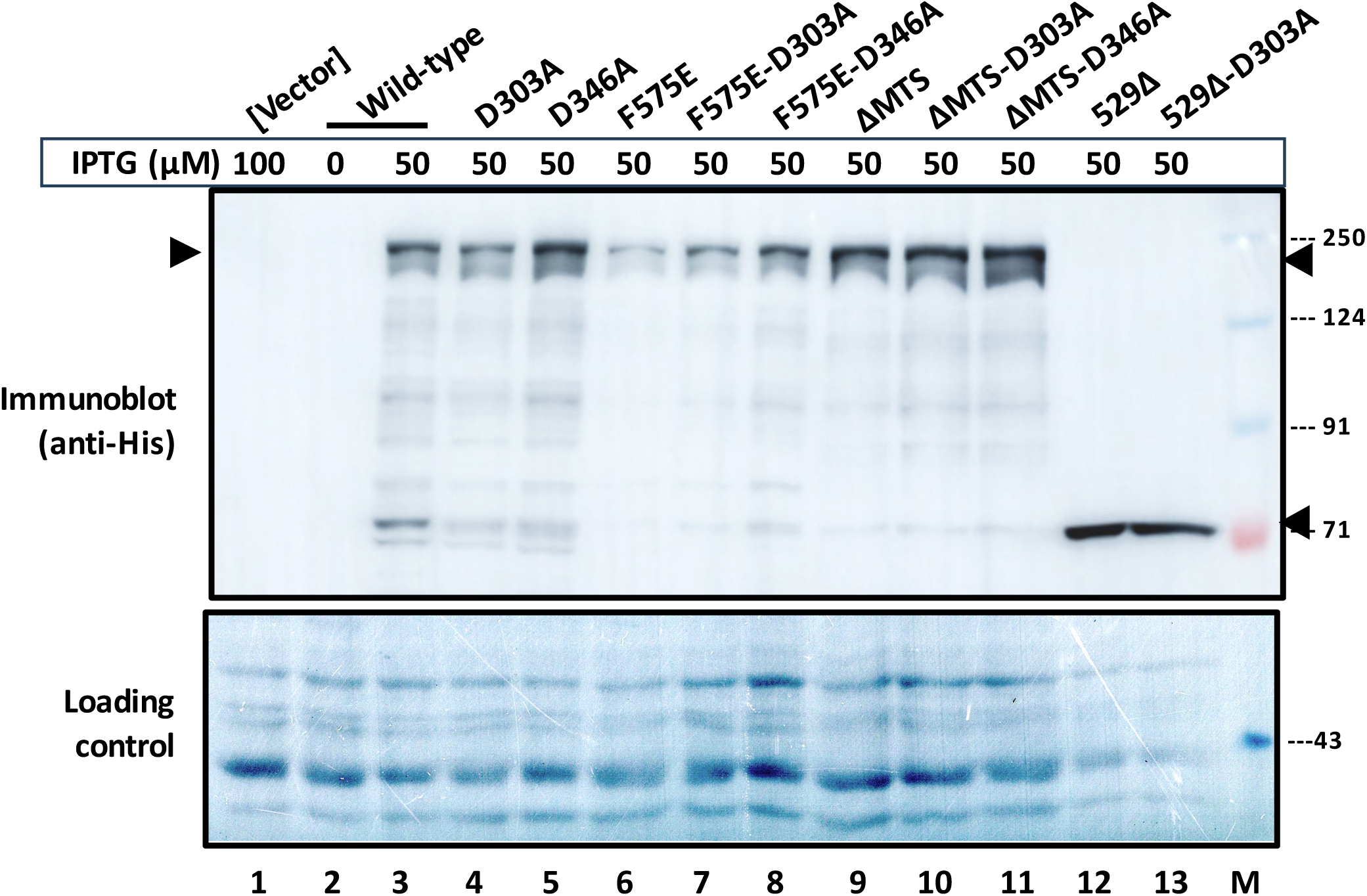
Determination by Western blotting of expression levels of wild-type RNase E and its variants in cultures of GJ15074 bearing ASKA plasmid vector or its derivatives encoding different RNase E polypeptides as indicated. IPTG supplementation of cultures was at the indicated concentrations. Top panel represents immunoblot with anti-His antibody, and the bottom panel represents the same blot stained with amido black (to serve as loading control). Amount of protein loaded on each of lanes 12-13 was one-half of that on each of lanes 1-11. Positions of migration of molecular-mass standards (in kDa) that were loaded on lane M are marked at right. Arrowheads denote the bands of wild-type RNase E and its variants.

**Supp. Fig. S4:**
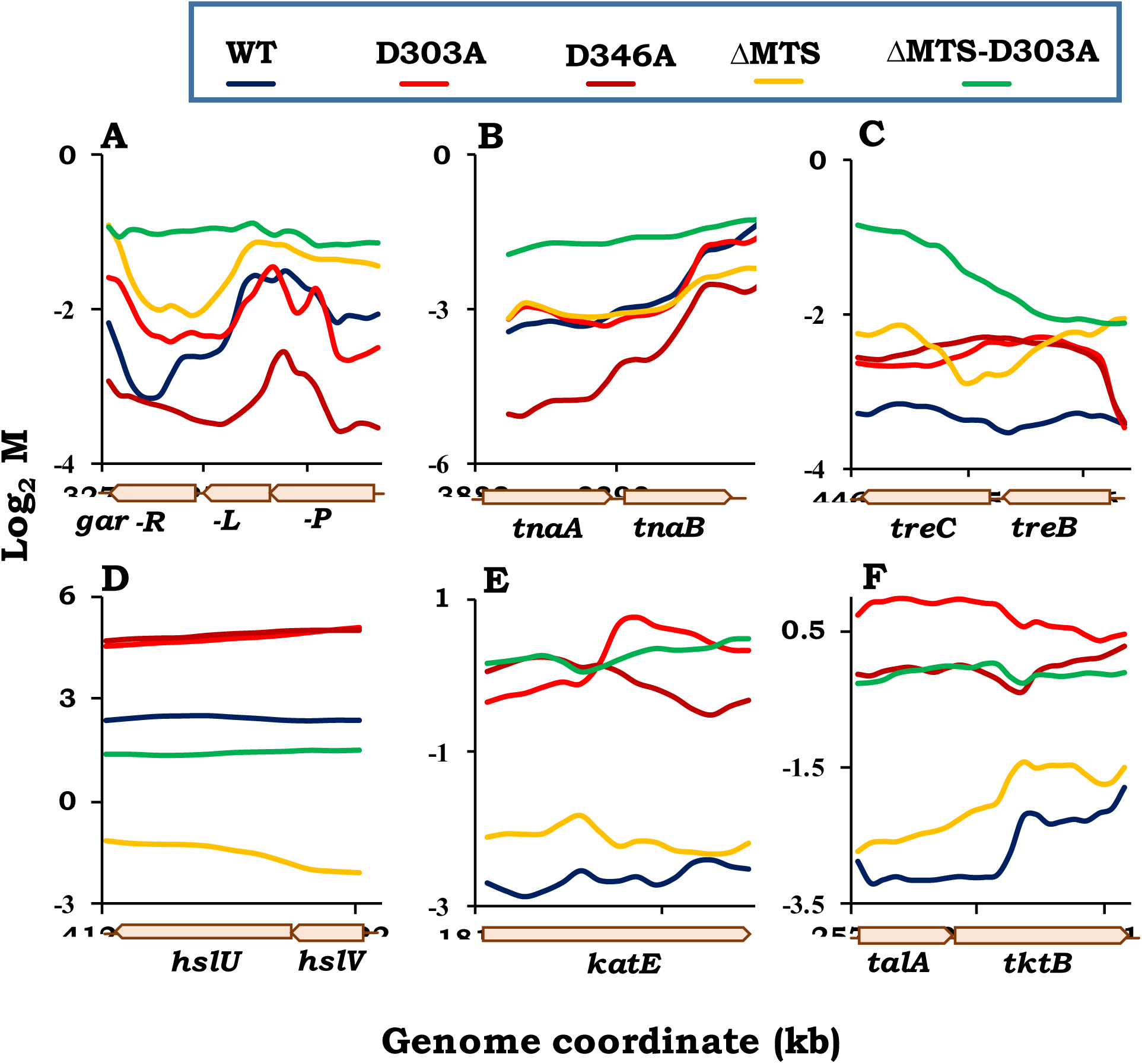
Plots of moving averages of log_2_ M values from RNA-Seq data at 100-base resolution for different genes in strains overexpressing wild-type (WT) RNase E or one of its variants (D303A, D346A, ΔMTS, and ΔMTS-D303A). Notations are as described in legend to Figure 4. Representative examples are shown of: (**A-C**) recessive resurrection in operons with negative log_2_ M; (**D**) genes of the σ^32^-heat shock regulon with positive log_2_ M; and (**E-F**) genes exhibiting discordant response upon overexpression of WT RNase E or its D303A/D346A variants.

**Supp. Fig. S5:**
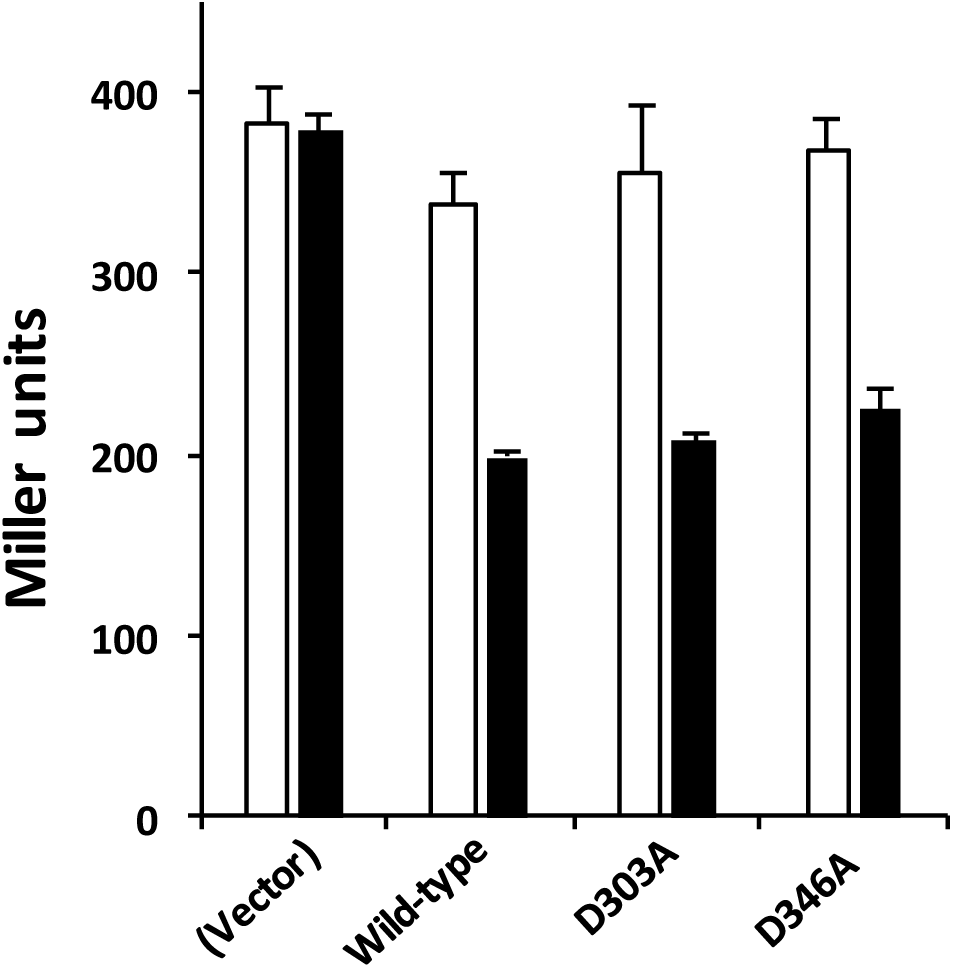
Specific activity (Miller units) of β-galactosidase in cultures without IPTG (open bars), or following supplementation with 100 μM IPTG for 30 min (filled bars), of *rne^+^ rne-lac* Δ*recA* strain GJ15074 carrying ASKA plasmid vector or its derivatives encoding different RNase E polypeptides as indicated. The decrease in *rne-lac* expression following IPTG supplementation was statistically significant for each of the derivatives with ASKA plasmids encoding wild-type RNase E as well as its D303A and D 346A variants (*P* = 0.0004, 0.0006 and 0.003, respectively; Student’s one-tailed *t*-test). Plasmids were (from left): pCA24N, ASKA-*rne*, pHYD5152 and pHYD5151.

**Supp. Fig. S6:**
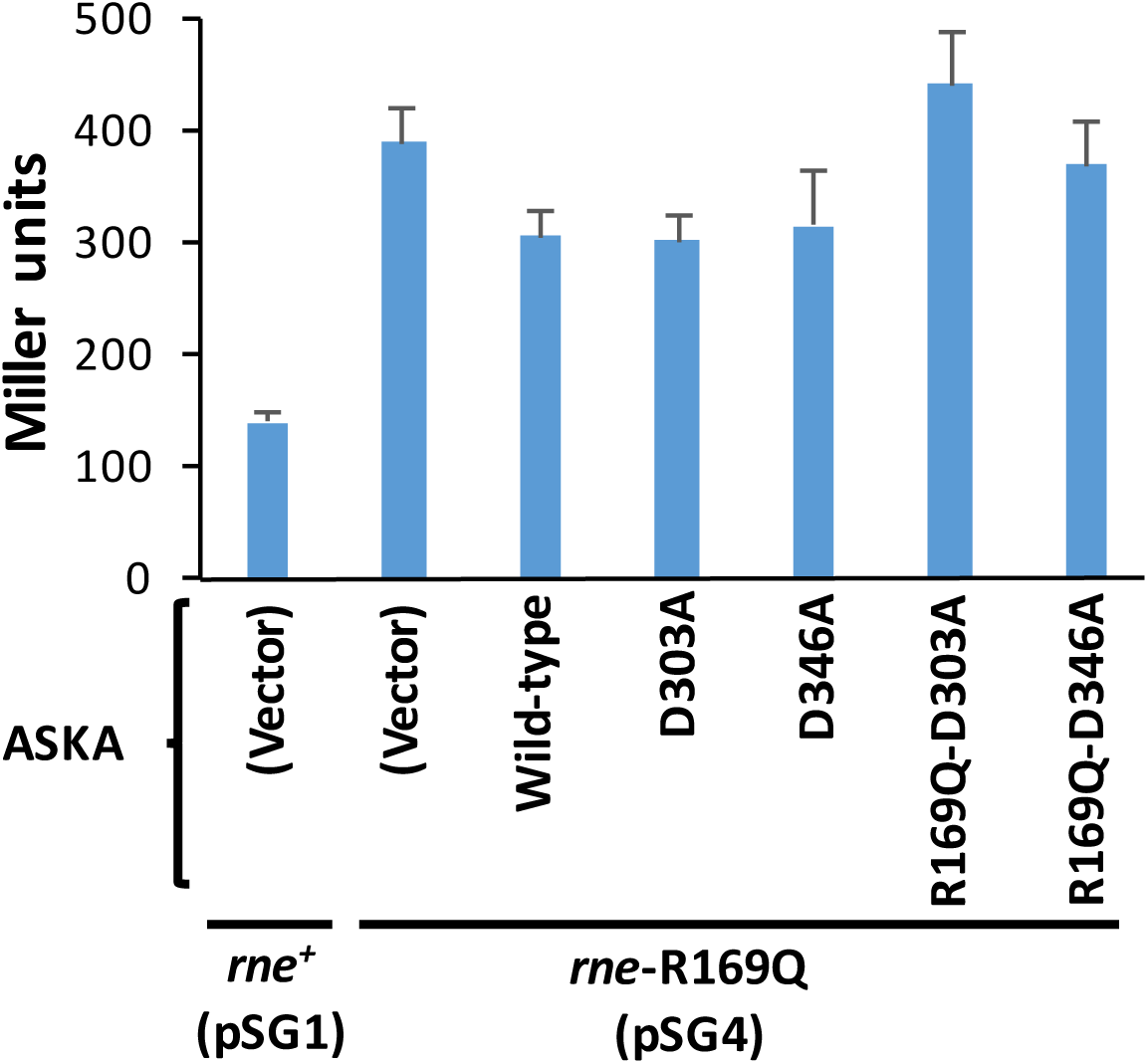
Specific activity (Miller units) of β-galactosidase in cultures supplemented with nil IPTG of derivatives of *rne-lac* Δ*recA* strains GJ20323/pSG1 (*rne^+^*) or GJ20323/pSG4 (*rne*-R169Q) carrying different plasmids as indicated. These values have been normalized to 1.0 for the different plots in Figure 5. Plasmids were (from left, numbers are to be prefixed with pHYD unless otherwise marked): pCA24N, pCA24N, ASKA-*rne*, 5152, 5151, 5165 and 5164.

